# PTHrP Induces STAT5 Activation, Secretory Differentiation and Mammary Tumor Progression

**DOI:** 10.1101/2021.12.28.473846

**Authors:** Diego Y. Grinman, Kata Boras-Granic, Farzin M. Takyar, Pamela Dann, Julie R. Hens, Christina Marmol, Jongwon Lee, Jungmin Choi, Lewis A. Chodosh, Martin E. Garcia Sola, John J. Wysolmerski

**Author notes:** Address Correspondence to: Diego Y. Grinman, PhD, Section of Endocrinology and Metabolism, Yale School of Medicine, 300 Cedar Street, TAC S120, Box 208020, New Haven, CT 06520-8020. Contributed Equally to this work (Diego Y. Grinman, Kata Boras-Granic, Farzin M. Takyar).

## Abstract

**Background:** Parathyroid hormone-related protein (PTHrP) is required for embryonic breast development and has important functions during lactation, when it is produced by alveolar epithelial cells and secreted into the maternal circulation to mobilize skeletal calcium used for milk production. PTHrP is also produced by breast cancers and GWAS studies suggest that it influences breast cancer risk. However, the exact functions of PTHrP in breast cancer biology remain unsettled.

**Methods:** We developed a tetracyline-regulated, MMTV (mouse mammary tumor virus)-driven model of PTHrP overexpression in mammary epithelial cells (Tet-PTHrP mice) and bred these mice with the MMTV-PyMT (polyoma middle tumor-antigen) breast cancer model to analyze the impact of PTHrP overexpression on normal mammary gland biology and in breast cancer progression.

**Results:** Overexpression of PTHrP in luminal epithelial cells caused alveolar hyperplasia and secretory differentiation of the mammary epithelium with milk production. This was accompanied by activation of Stat5 and increased expression of E74-like factor-5 (Elf5). In MMTV-PyMT mice, overexpression of PTHrP (Tet-PTHrP;PyMT mice) shortened tumor latency and accelerated tumor growth, ultimately reducing overall survival. Tumors overproducing PTHrP also displayed increased expression of nuclear pSTAT5 and Elf5, increased expression of markers of secretory differentiation and milk constituents, and histologically resembled secretory carcinomas of the breast. Overexpression of PTHrP within cells isolated from tumors, but not PTHrP exogenously added to cell culture media, led to activation of STAT5 and milk protein gene expression. In addition, neither ablating the Type 1 PTH/PTHrP receptor (PTH1R) in epithelial cells or treating Tet-PTHrP;PyMT mice with an anti-PTH1R antibody prevented secretory differentiation or altered tumor latency. These data suggest that PTHrP acts in a cell-autonomous, intracrine manner. Finally, expression of PTHrP in human breast cancers is associated with expression of genes involved in milk production and STAT5 signaling.

**Conclusions:** Our study suggests that PTHrP promotes pathways leading to secretory differentiation and proliferation in both normal mammary epithelial cells and in breast tumor cells.

## Background

Parathyroid hormone–related protein (PTHrP) was originally discovered as a cause of elevated calcium levels in patients with cancer [1–3]. It is evolutionarily related to parathyroid hormone (PTH) and the amino-terminal portions of both proteins are highly homologous, allowing them to bind and activate the same Type 1 PTH/PTHrP receptor (PTH1R) [2, 3]. As a result, when PTHrP is secreted by tumors, it mimics PTH, leading to excessive bone resorption and hypercalcemia. PTHrP also contributes to the development and physiologic functions of a variety of tissues and it has been shown to affect cell proliferation and cell death in a number of settings [2, 4–6]. While many of its functions are mediated by the PTH1R, PTHrP can also remain within the cell to regulate proliferation, differentiation and survival through an intracrine mode of action requiring the translocation of PTHrP into the nucleus [3, 7–11]. Although nuclear translocation appears to be important for PTHrP biology, details of this signaling pathway remain obscure.

PTHrP and the PTH1R are expressed throughout the life cycle of the mammary gland as well as in breast tumors. Both molecules are required for fetal breast development in mice and humans [12–15]. PTHrP also has important functions during lactation. Its production is greatly upregulated in alveolar epithelial cells and it is secreted into both milk and the maternal circulation [16–18]. In the maternal circulation, PTHrP acts on bone cells to mobilize calcium from the skeleton that is subsequently used by the mammary gland for milk production. In addition, PTHrP in milk regulates total body calcium accrual in suckling neonates, acting to coordinate maternal and neonatal calcium economy [19].

PTHrP is also produced by breast cancers, contributing both to their growth and to tumor-induced changes in systemic metabolism [6, 16, 20]. When produced by breast cancer cells within the bone microenvironment, PTHrP contributes to osteolytic bone destruction and the expansion of bone metastases [6, 21, 22]. In addition, genome-wide association (GWAS) studies have implicated the *PTHLH* (PTHrP) gene as a breast cancer susceptibility locus [16, 23–25], suggesting that it may contribute to early steps in transformation and/or cancer progression. However, the exact functions of PTHrP in breast cancer biology remain unsettled. Different studies have reported that its expression either correlates with increased or decreased metastases and survival [11, 26–30]. Moreover, studies have variably reported that PTHrP either stimulates or inhibits the proliferation, differentiation and survival of breast cancer cells [5, 6, 11, 22, 31–34]. These contradictory results concerning the role and prognostic value of PTHrP expression in breast cancer underscore the need to better understand how it modulates breast tumor growth and/or breast cancer susceptibility.

In order to examine the effects of PTHrP on mammary tumor development in mice, we developed a tetracyline-regulated, MMTV-driven model of PTHrP overexpression in mammary epithelial cells (MMTV-rtTA;tetO-PTHrP). We found that overexpression of PTHrP in luminal epithelial cells caused alveolar hyperplasia and secretory differentiation of the mammary epithelium enabling virgin mice to produce milk. This phenotype was associated with activation of STAT5, and increased expression of Elf5 (E74-like factor-5), both important regulators of alveolar secretory differentiation [35–38]. Furthermore, overexpression of PTHrP in epithelial cells in MMTV-PyMT mice dramatically promoted the formation of mammary tumors by shortening tumor latency and accelerating tumor growth, ultimately reducing overall survival. Interestingly, tumors overproducing PTHrP expressed markers of secretory differentiation and expressed milk constituents. These data suggest that PTHrP promotes pathways leading to secretory differentiation in both normal mammary epithelial cells and in breast tumor cells.

## Methods

### Animals

We used FVB female mice of various genotypes described below in all experiments. Male mice were not used because the focus of the study was on mammary gland development and breast cancer. All animal experiments were performed in accordance with institutional regulations after protocol review and approval by Yale University’s Institutional Animal Care and Use Committee.

Six different genetically engineered mouse models were used in this study: MMTV-rtTA, MMTV-rtTA;TetO-PTHrP, MMTV-PyMT, MMTV-rtTA;TetO-PTHrP;MMTV-PyMT, MMTV-rtTA; TetO-PTHrP;MMTV-PyMT;MMTV-Cre and, MMTV-rtTA;TetO-PTHrP;MMTV-PyMT; MMTV-Cre;PTH1R^lox/lox^. We used a bi-transgenic, tetracycline-regulated, mouse mammary tumor virus long terminal repeat (MMTV) system to control the timing of PTHrP overexpression. MMTV-rtTA mice from the Chodosh laboratory (University of Pennsylvania) [39] were bred to TetO-PTHrP responder mice generated in by the Wysolmerski laboratory [40] to make the double transgenic MMTV-rtTA;TetO-PTHrP (Tet-PTHrP) mice. MMTV-PyMT mice were purchased from Jackson Laboratories on a FVB background and bred into our Tet-PTHrP mice to generate MMTV-rtTA;TetO-PTHrP;MMTV-PyMT (Tet-PTHrP;PyMT) mice. MMTV-Cre (Jackson Laboratories) and PTH1R^fl/fl^ mice (from Henry Kronenberg, Boston, MA) [41] were bred into the MMTV-rtTA;TetO-PTHrP;MMTV-PyMT mice to generate MMTV-rtTA;TetO-PTHrP;MMTV-PyMT;PTH1R^lox/lox^ (Tet-PTHrP;PyMT;PTH1RLox) and MMTV-rtTA;TetO-PTHrP;MMTV-PyMT;MMTV-Cre;PTH1R^lox/lox^ (Tet-PTHrP;PyMT;Cre;PTH1RLox) mice.

Doxycycline (Dox) (2mg/ml; Research Products International, Cat# D43020) was administered in 5% sucrose water and mice could drink ad libitum. Mice were followed weekly for tumors. Once palpable, tumor size was measured weekly with calipers and tumor volume calculated using the formula 0.5 x length x width^2^. Mice were euthanized when tumors reached approximately 1.5 cm in any dimension, or when they appeared unhealthy during the course of the experiment, whichever was earlier.

### Biochemical measurements

Serum calcium concentrations were measured using the Quantichrom Calcium Assay Kit (DICA-500, BioAssay Systems) according to manufacturer’s instructions. Plasma PTHrP was measured using an immunoradiometric assay (DSL-8100; Beckman Coulter) in which we substituted a rabbit anti-PTHrP (1–36) antibody generated in our laboratory as capture antibody [42]. This assay has a sensitivity of 0.3 pM.

### Whole-mount analysis

Whole-mount analysis was performed on mammary tissue as previously described [43]. Briefly, the no. 4 inguinal mammary glands were removed and mounted on a microscope slide. The tissue was fixed in acid ethanol for 1 h at room temperature, washed in 70% ethanol then distilled water and incubated in carmine aluminum stain (0.2% carmine, 0.5% aluminum potassium sulfate) overnight at room temperature. After staining, the mammary glands were dehydrated through graded ethanol and cleared in acetone and then toluene before being mounted under glass coverslips using Permount (Fisher Scientific, Cat# SP15-100).

### Histology and immunohistochemistry

Two hours prior to euthanasia, mice were injected with BrdU (Roche) or EdU (50mg/kg, Invitrogen). Whole mammary glands, tumors and lungs were removed, weighed and fixed for 12 hours in 4% paraformaldehyde. After fixation in 4% paraformaldehyde, tissues were transferred to 70% ethanol, embedded in paraffin and cut in 5 μm thick sections. Pertinent slides were then either stained with hematoxylin and eosin using standard conditions, used for immunohistochemistry, or processed for measuring proliferation using anti-Bromodeoxyuridine-POD, Fab fragment Kit (Roche, Cat# 11585860001) or the Click-iT EdU Cell Proliferation Kit (Invitrogen Cat# C10337). Rates of proliferation were calculated by dividing the number of BrdU- or EdU-positive nuclei by the total number of nuclei. Lungs were processed for histology and pulmonary metastases quantified by examination of 10, H&E-stained sections cut 105 µm apart. All immunohistochemistry included an IgG isotype control and the primary antibodies we used were against phospho-Stat5 (Cell Signaling, Cat# 9314), β-casein (Santa Cruz Biotechnology, Cat# sc-166530), Elf-5 (Santa Cruz Biotechnology, Cat# sc-9645), NF1B (Sigma, Cat# HPA-0039556), Nkcc1 (gift from Dr. James Turner at National Institutes of Health) and Npt2b (gift from Dr. Jürg Biber at University of Zurich). Staining was detected using Vector Elite ABC kits (Vector Laboratories), Envision Plus (DAKO), or M.O.M. Immunodetection Kit (Vector Laboratories, Cat) and we used 3,3′-diaminobenzidine as a chromogen.

### RNA extraction and real-time RT-PCR

Mammary glands and tumors were homogenized in 1 ml TRIzol (Invitrogen, Cat# 15596018) using an Ultraturrax T25 (Ika Labortechnik) on ice. Lysates were cleared at 13,000 *g* for 10 min at 4°C. The RNA was isolated using PureLink RNA columns (Invitrogen, Cat# 12183025) according to the manufacturer’s instructions. Total RNA was quantified using a Nanodrop 1000 spectrophotometer (Thermo Fisher Scientific). For all samples, the ratio of absorbance at 260 nm to absorbance at 280 nm was >1.8. cDNA was synthesized using 1 μg of total RNA with the High Capacity cDNA Reverse Transcription Kit (Applied Biosystems, Cat# 4368814) according to the manufacturer instructions. Quantitative RT-PCR was performed using the Taqman Fast Universal PCR Master Mix (Applied Biosystems, Cat# 4352042) or Sybr Green PCR Master kit (Applied Biosystems, Cat# 4309155) and a StepOnePlus real-time PCR system (Applied Biosystems). The following TaqMan primer sets were used: hPthlh Hs00174969_m1, mPthlh Mm00436057, PTH1R Mm00441046, Wap Mm00839913_m1, Lalba Mm00495258_m1, Csn1s1 Mm01160593_m1, Csn1s2a Mm00839343_m1, Csn1s2b Mm00839674_m1, Csn2 Mm04207885_m1, Csn3 Mm02581554_m1, Elf5 Mm00468732_m1, Nfib Mm01257777_m1, Gata3 Mm01337570_m1, Hprt1 Mm03024075_m1, Actb (Cat# 4352933E). The following primer pairs for Sybr green were also used: Pymt fwd (5′-ctgctactgcacccagacaa-3′) and Pymt rev (5′-gcaggtaagaggcattctgc-3′), Actb fwd (5’-ccacacccgccaccagttc-3’) and Actb rev (5’-gacccattcccaccatcacacc-3’). Relative mRNA expression was determined using the standard curve method with the StepOne software v2.3 (Applied Biosystems).

### Tissue protein isolation and western blot

Pieces of mammary gland or mammary tumor no more than 0.5 cm x 0.5 cm were lysed in 1ml of RIPA lysis buffer (10 mM Tris·HCl pH 8, 140 mM NaCl, 1 mM EDTA pH 8, 0.5mM EGTA pH 8, 1% Triton X-100, 0.1% deoxycholate, and 0.1% SDS) supplemented with a cocktail of protease inhibitors (Thermo Scientific Cat# 78429), 50 mM NaF, and 1 mM Na3VO4 on ice. Samples were then homogenized using an Ultraturrax T25 (Ika Labortechnik). Lysates were centrifugated at 13,000 g for 10 min at 4°C and the supernatant was recovered. The samples were quantitated for total protein using the Bradford protein assay (Bio-Rad Cat# 5000001) following the manufacturer’s instructions. A 2µg/µl protein solution containing sample buffer (Invitrogen Cat# NP0007) plus sample reducing agent (Invitrogen Cat# NP0004) was prepared and 30ug of total protein were loaded into precast, 4% to 12% Bis-Tris acrylamide gels (Thermo Fisher Scientific, Cat# NP0322) in MOPS buffer (Thermo Fisher Scientific, Cat# NP0001) and underwent electrophoresis, after which samples were transferred to nitrocellulose membranes (Bio-Rad, Cat# 1621112). Membranes were treated with blocking buffer (LI-COR Biosciences, Cat# 927-60001) for 1 hour at room temperature and then incubated with the primary antibody overnight at 4°C, followed by a dye conjugated secondary antibody for 1 hour at room temperature. Membranes were imaged and analyzed using the Odyssey IR imaging system (LI-COR Biosciences). The primary antibodies used were: anti-PTHrP (Peprotech, Cat# 500-P276), anti-β-casein (Santa Cruz Cat# 166530), anti-Elf5 (Santa Cruz Cat# sc-9645), anti-NF1B (Sigma, Cat# HPA003956), anti-p(Tyr694)Stat5 (Cell Signaling, Cat# #9359), anti-Stat5 (Cell Signaling, Cat# #94205), anti-Npt2b (gift from Dr. Jürg Biber at University of Zurich), anti-β-Actin (Santa Cruz Cat# sc-130656). The secondary antibodies used were anti-mouse (LI-COR, Cat# 926-68022) and anti-Rabbit (LI-COR Biosciences, Cat# 926-32213)

### Tumor cell isolation and culture

Tumor cells were isolated from transgenic mammary tumors as previously described [42]. Briefly, dissected tumors were minced into fragments under sterile conditions and subjected to enzymatic digestion with Collagenase-Type3 (Worthington, Cat#: LS004183) at 2mg/ml, Dispase (Gibco, Cat#: 17105-041) at 2mg/ml, Gentamycin (Gibco, Cat#:15710-064) at 50µg/ml, Amphotericin B (Sigma, Cat#:A2942) at 250µg/ml, and 5% FBS in DMEM/F12 media for 3 hours with intermittent shaking. Following digestion, tumor organoids were pelleted, and then treated with NH4Cl (Stem Cell Technologies, Cat # 07800) to lyse RBCs, following which, the pellet was washed three times with PBS. Organoids were then passed through a 70 µm cell strainer, counted and used for transplantation experiments or cultured at a density of 3×10^6^ cells/55cm^2^. Proliferation of cultured cells was measured by assessing BrdU incorporation (Cell proliferation ELISA Kit 11647229001; Roche) after addition of Dox (2µg/ml) or PTHrP (Bachem, Cat# 4017147.0500) to the culture media.

### Tumor cell transplantation

500,000 freshly isolated, sterile tumor cells were suspended in 150 µl of sterile saline and were injected subcutaneously into the fat pad of 8 wild-type, adult FVB mice via a small incision between the fourth nipple and the midline as previously described [44]. Mice were treated with Dox 24 hours prior to the injection and were monitored for tumor development. Mice were checked twice a week for tumors and tumor size was measured with calipers every other day. Tumor-bearing animals were euthanized when the tumors reached approximately 1.5 cm in any dimension or when they appeared unhealthy, whichever was earlier.

### Global gene expression profiling

Total RNA was prepared using TRIzol reagent (Invitrogen) from FACS sorted luminal epithelial cells of 4.5 week-old, MMTV-rtTA and Tet-PTHrP mice on Dox from birth using antibodies against CD24 and CD49f cell surface markers as previously described [45]. Similarly, total RNA was prepared from whole tumor lysates of MMTV-PyMT and Tet-PTHrP;PyMT mice on Dox. The isolated RNA was purified using the RNeasy cleanup kit (Qiagen). RNA was reverse-transcribed and hybridized to Affymetrix Mouse Genome 430 2.0 GeneChip by the Yale Center for Genomic Analysis. Microarray data were analyzed with R version 4.1.2 and Bioconductor 3.14 [46]. Raw data were MAS5 normalized and log_2_ transformed. 20,000 probes with the highest statistical significance were selected as the first working matrix and then only genes with fold change of +/− 2 and p<0.01 were considered for further analyses. Differentially expressed genes (DEGs) were analyzed using WikiPathways Pathway Analysis for biological interpretation [47] and significant pathways were based on the Bonferroni adjusted p value (padj) <0.05. Results of the functional analysis were combined and integrated to the expression data with the GOplot package [48]. All statistical analyses and data visualization plots were done with R/Bioconductor packages. GSEA analysis was performed using previously generated set of ∼200 STAT5-dependent and mammary tissue restricted genes [38]. Enrichment score curves and member ranks were generated by the GSEA software package [49]. Volcano plots were constructed from the first selected 20,000 probes matrix with *ggplot2* [50]. Heatmap was generated with *heatmap.2* package.

### Breast cancer single cell RNA seq data download and process

Count matrices from published single cell RNA sequencing (scRNA-seq) datasets were downloaded from the NCBI Gene Expression Omnibus (GSE161529) and then analyzed using Seurat version 4.0 [51]. Seurat objects were created from 15 ER+, 6 HER2+ and 4 TNBC patient-derived datasets. Cells with > 60,000 counts and the number of unique genes detected in each cell were removed using > 200 and < 7,000 as criteria. This is a quality control step, as it is thought that cells with high numbers of counts are more likely to be doublets while cells with low numbers of counts are thought to be of poor data quality. Data normalization, variable feature detection, feature scaling, and principal component analysis were performed in Seurat using default parameters. Cell clusters were identified using the default Louvain clustering algorithm implemented in Seurat. Default Seurat function settings were used except that clustering resolutions were set to 0.5 and principal component dimensions 1:10 were used for all dimension reduction and integration steps. Epithelial cells were identified using canonical marker genes as described and normalized counts data were used in all relevant downstream analysis [52]. Cells were divided into two groups depending on their normalized counts of *PTHLH* expression level. *PTHLH* high groups expressed *PTHLH* more than 0 and remaining cells were designated as the *PTHLH* low group. Differential expression between PTHLH high and low groups was conducted using the FindMarker function in Seurat package with MAST option. Pathway enrichment was performed on ranked lists with fGSEA using HALLMARK gene set from MsigDB v7.4 [49, 53]. After removing genes that are not expressed in any cell, protein coding genes only were considered (refer to *biomaRt* package [54]).

### Statistics

Results were expressed as means ± SE of at least 3 independent experiments. Statistical analyses were performed with Prism 9.0 (GraphPad Software) and consisted of one-way ANOVA followed by Tukey’s multiple comparisons test. Before statistical analysis, Q-Q plot and Shapiro Wilks test were performed for normality. Homoscedasticity was assessed with Levene’s test. In figures, asterisks mean significant differences between means.

## Results

### Overexpression of PTHrP in luminal epithelial cells causes alveolar hyperplasia

We created a tetracycline-regulated model of PTHrP overexpression using a well described MMTV-rtTA mouse, that employs the mouse mammary tumor virus long terminal repeat (MMTV) to drive expression of the reverse tetracycline transactivator (rtTA) in mammary epithelial cells (MECs) [55]. When MMTV-rtTA mice were bred to a mouse containing a tetracycline-responsive, human PTHrP transgene, (TetO-PTHrP mice) [56], the resulting double-transgenic, MMTV-rtTA;TetO-PTHrP (Tet-PTHrP) offspring demonstrated a significant induction of human *PTHLH* mRNA in mammary glands upon treatment with Dox (Fig.1A). As expected, there was essentially no human *PTHLH* mRNA expressed in mammary glands from Tet-PTHrP mice in the absence of Dox, nor was there induction of the endogenous mouse *Pthlh* gene in response to Dox.

**Figure 1.**
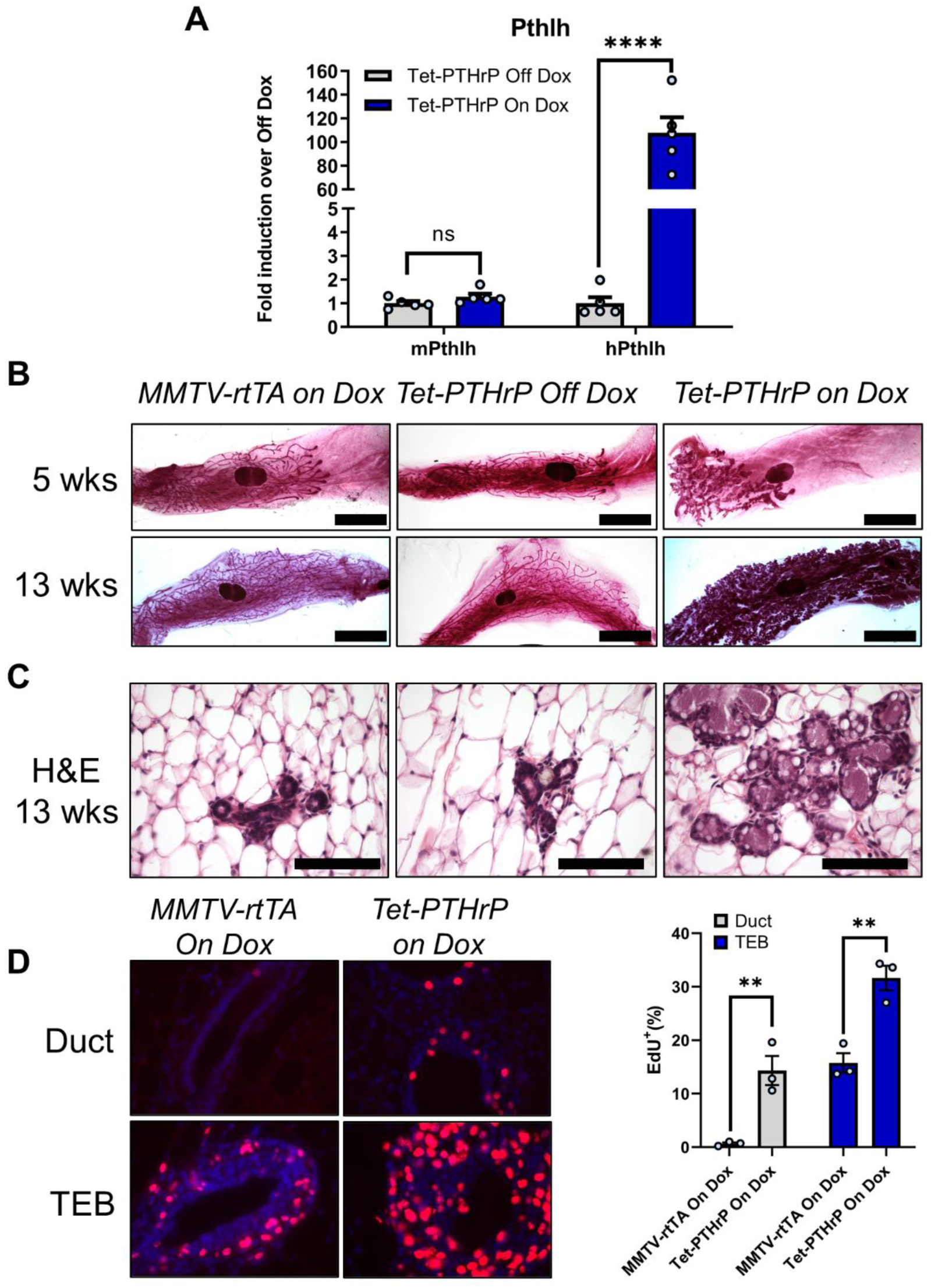
PTHrP overexpression causes alveolar hyperplasia. A) Relative expression of mouse and human *PTHLH* mRNA in mammary gland lysates. *Hprt1* was used as housekeeping gene. Bars represent mean ± SEM of fold change versus Tet-PTHrP off dox, n=5 mice per group. B) Whole-mount analysis of carmine-stained number 4 inguinal mammary glands from mice at 5 and 13 weeks. Scale bar 5mm. C) H&E stained cross-sections from 13-week-old mice. Scale bar 100µm. D) Representative images and quantification of EdU incorporation in sections of mammary glands from mice at 5 weeks of age. Bar graphs represent the percentage of Edu-positive cells over a minimum of 1000 total nuclei (DAPI). N=3 mice per group, ****p<0.0001 **p<0.01.

In previous studies, overexpression of PTHrP in myoepithelial cells delayed ductal elongation and this was also the case in Tet-PTHrP mice treated with Dox [56]. As shown in Fig.1B, at 5 weeks of age, the ducts in Tet-PTHrP virgin mice off Dox had grown past the central lymph and displayed a dichotomous branching pattern typical of the virgin mammary gland. By contrast, in Tet-PTHrP mice treated with Dox, the ducts were foreshortened but hyperplastic in appearance. Ducts in MMTV-rtTA mice on Dox were the same as those in Tet-PTHrP mice off Dox, demonstrating that the effects observed were due to PTHrP and not to either Dox or rtTA. By 13 weeks of age, the ducts in all genotypes had advanced to the borders of the fat pad but the glands of Dox-treated Tet-PTHrP mice displayed obvious alveolar hyperplasia on whole mount, reminiscent of normal glands in mid to late pregnancy. In fact, histologic examination of mammary glands from the Dox-treated Tet-PTHrP mice demonstrated multiple clusters of MECs forming alveolar structures (Fig. 1C). Since there was no apparent structural or developmental difference between MMTV-rtTA mice on Dox and Tet-PTHrP mice off Dox, nor leakiness of human PTHrP expression in the latter, subsequent experiments used either of these genotypes interchangeably as controls.

Given the increased numbers of epithelial cells in the glands from Tet-PTHrP mice treated with Dox, we assessed rates of epithelial cell proliferation by measuring EdU incorporation. As shown in Fig 1D, there was a clear increase in EdU incorporation in MECs in both ducts and terminal end buds (TEBs) in response to PTHrP overexpression. These data demonstrate that induction of PTHrP expression in MECs leads to alveolar hyperplasia.

### Overexpression of PTHrP Activates Secretory Differentiation of Mammary Epithelial Cells

Overexpression of PTHrP was accompanied by distension of the hyperplastic alveolar and ductal lumens with what appeared to be secretory material. In addition, large lipid droplets were apparent in both the cells as well as the luminal space (Fig. 1C). These features suggested milk production, and milk-like fluid was evident upon gross inspection of the intact mammary glands of Tet-PTHrP mice treated with Dox (Additional File 1A).

In order to confirm that MECs underwent secretory differentiation in response to PTHrP, we assayed differentiation markers typically expressed by MECs during lactation [35, 57]. As shown in Fig. 2A, Dox treatment induced the expression of β-casein and the sodium-phosphate transporter 2b (NPT2b) as measured by immunohistochemistry, while suppressing expression of the sodium-potassium-chloride co-transporter (NKCC1) in MECs of virgin Tet-PTHrP mice. These changes were identical to normal lactating controls but were absent in normal virgin controls and in virgin Tet-PTHrP mice in the absence of Dox. We also assayed the expression of a series of milk-protein genes by QPCR (Fig 2B). Overexpression of PTHrP caused the induction of whey acidic protein (*Wap*), alpha lactalbumin (*Lalba*) and multiple casein genes, none of which were expressed in the glands from virgin controls or in Tet-PTHrP mice off Dox. As expected from the gene expression data and the immunostaining, PTHrP overexpression led to a significant increase in PTHrP protein levels as well as an increase in β-casein and NPT2b protein expression in whole mammary gland lysates as assessed by immunoblot (Fig 2C & Additional File 1B).

**Figure 2.**
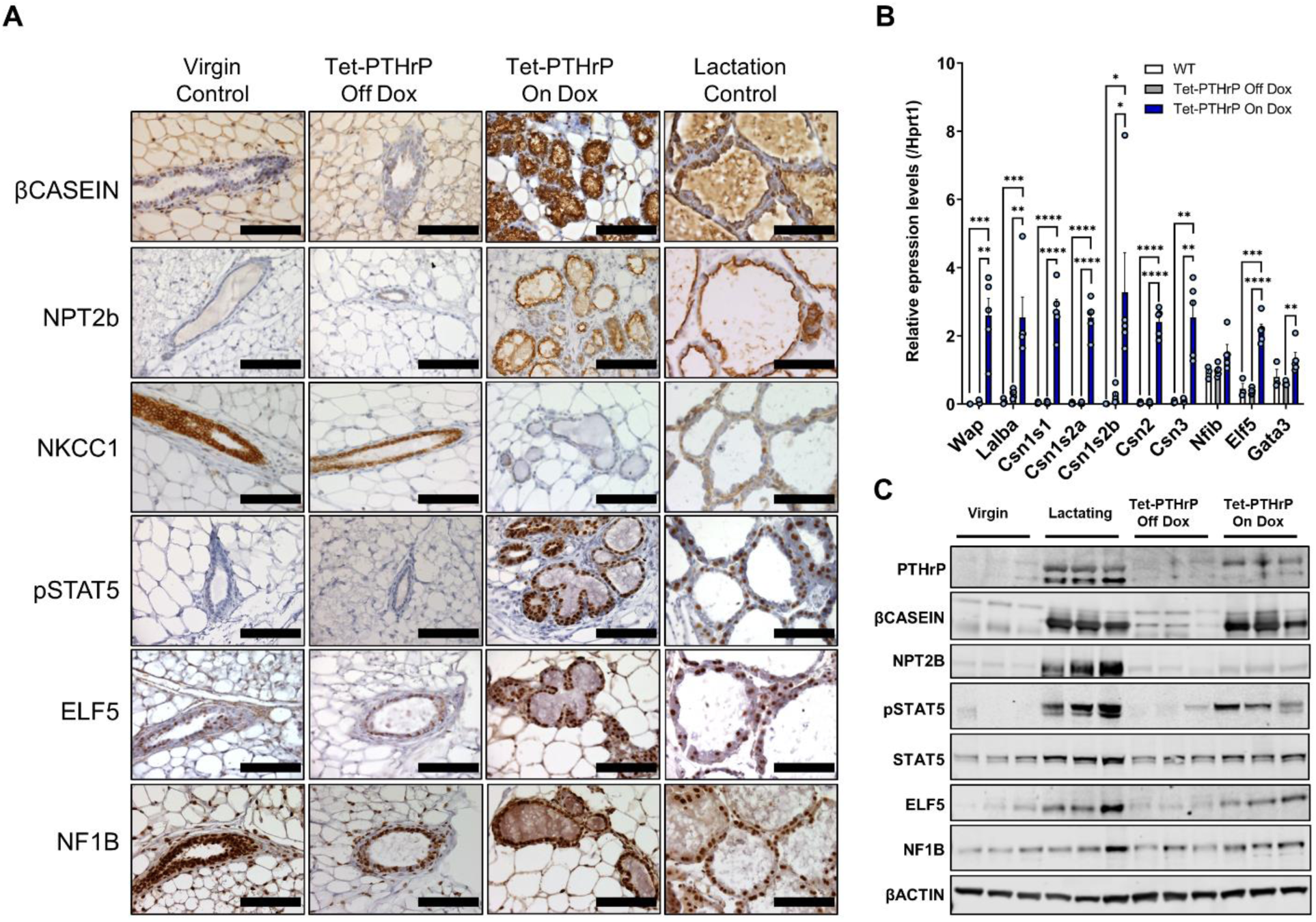
PTHrP overexpression induces the expression of milk proteins and markers of secretory differentiation. A) Immunohistochemical staining of mammary gland sections of 8-12-week-old mice. Representative images from each group are shown. N=3, Scale bar 100 µm. B) QPCR analysis for the expression of milk protein and transcription factor genes from whole mammary glands of 8-12-week-old mice. *Hprt1* was used as housekeeping gene. C) Western blot analysis of protein lysates from mammary glands of 8-12-week-old mice. Samples from three different mice per group were run with β-Actin as the loading control. Bars represent mean ± SEM, n=3 per group, ****p<0.0001 ***p<0.001 **p<0.01 *p<0.05.

Next, we examined whether alveolar hyperplasia and/or secretory differentiation in MECs required ongoing exposure to PTHrP. Female Tet-PTHrP mice were placed on Dox at 8 weeks of age for 4 weeks and then were either euthanized immediately or followed for an additional 6 weeks off Dox before being euthanized. Additional controls included similar nulliparous, Tet-PTHrP mice treated with Dox for 10 continuous weeks before euthanasia. As expected, PTHrP expression for 4 weeks or 10 weeks in adult females triggered alveolar hyperplasia, activated the expression of β-casein and NPT2b, and suppressed NKCC1 expression (Additional File 2). After the withdrawal of Dox for 6 weeks, the alveolar hyperplasia had almost completely resolved histologically, NPT2b staining was no longer observed and NKCC1 became evident. However, MECs continued to express some β-casein, albeit principally within the lumen of the ducts and at lower levels than in MECs from glands exposed to ongoing PTHrP overexpression. These results suggest that alveolar hyperplasia and the mature secretory phenotype requires ongoing exposure to PTHrP but that some changes in cell differentiation or cell fate may persist after transient exposure to PTHrP.

### Overexpression of PTHrP induces Changes in Gene Expression Similar to Lactation

Secretory differentiation of MECs during normal pregnancy and lactation requires changes in gene expression driven by several pioneering transcription factors including pSTAT5, Elf 5 and Nuclear factor 1B (NF1B) [35, 38, 58, 59]. As seen in Fig. 2A, immunohistochemistry demonstrated that Dox treatment of virgin Tet-PTHrP mice induced the expression of pSTAT5 within MEC nuclei, mimicking the pattern typically seen during lactation. There was also a clear increase in pSTAT5 in immunoblot analyses of mammary glands taken from Tet-PTHrP mice on Dox, that was not present in Tet-PTHrP mice off Dox (Fig. 2C and Additional File 1B). While nuclear staining for ELF5 was evident in MECs from virgin controls and from Tet-PTHrP mice off Dox, the staining intensity appeared increased in MECs from virgin Tet-PTHrP mice on Dox and in lactating control mice (Fig. 2A). This impression was confirmed by an increase in *Elf5* mRNA as assessed by QPCR (Fig 2B) as well as increased ELF5 protein levels as measured by immunoblot (Fig 2C and Additional File 1B). Finally, expression of NF1B as assessed by immunohistochemistry, QPCR and immunoblot was not clearly different in lactating mammary glands or in mammary glands from Tet-PTHrP mice on Dox as compared with glands from either control Tet-PTHrP mice off Dox or virgin mice. Continued full expression of these transcription factors required the ongoing presence of PTHrP, because withdrawal of PTHrP expression resulted in substantial reduction, although not complete elimination of the immunostaining for pSTAT5 and ELF5. As before, expression of NF1B was not affected by PTHrP expression (Supplemental Fig. 2).

We next performed an analysis of overall gene expression using oligonucleotide-based microarrays. We compared mRNA expression patterns from luminal MECs isolated from Tet-PTHrP mice on Dox to that of luminal MECs isolated from MMTV-rtTA control mice on Dox. Using a log fold change (LFC) cutoff of 2 and an adjusted p value of 0.01, we found 1631 genes differentially expressed (597 increased and 1034 reduced) as a result of PTHrP expression (Fig. 3A). Pathway analysis demonstrated that the differentially expressed transcripts comprised key pathways important for MEC secretory differentiation, including PI3K/Akt signaling, fatty acid biosynthesis, triglyceride biosynthesis and the mammary gland transition from pregnancy to lactation (although this didn’t quite reach statistical significance, padj=0.07), among others (Fig. 3B). A more detailed analysis of the genes involved in alveolar cell differentiation revealed an increase in the levels of *Elf5, Nf1b, Gata3, Sox9, Csn3, Tfap2c, PiK3r1, Lalba, Cldn8* and Muc1 as well as a downregulation of *Esr1, Pgr, Cav1, Cdo1* and *Ccnd2* transcripts, all changes consistent with the activation of a secretory program in luminal cells similar to lactation [60–62] (Fig 3C). Taken together, these results demonstrate that increased levels of PTHrP are sufficient to induce the expression of lactation-associated transcription factors that subsequently cause secretory differentiation and milk production in the absence of a prior pregnancy.

**Figure 3.**
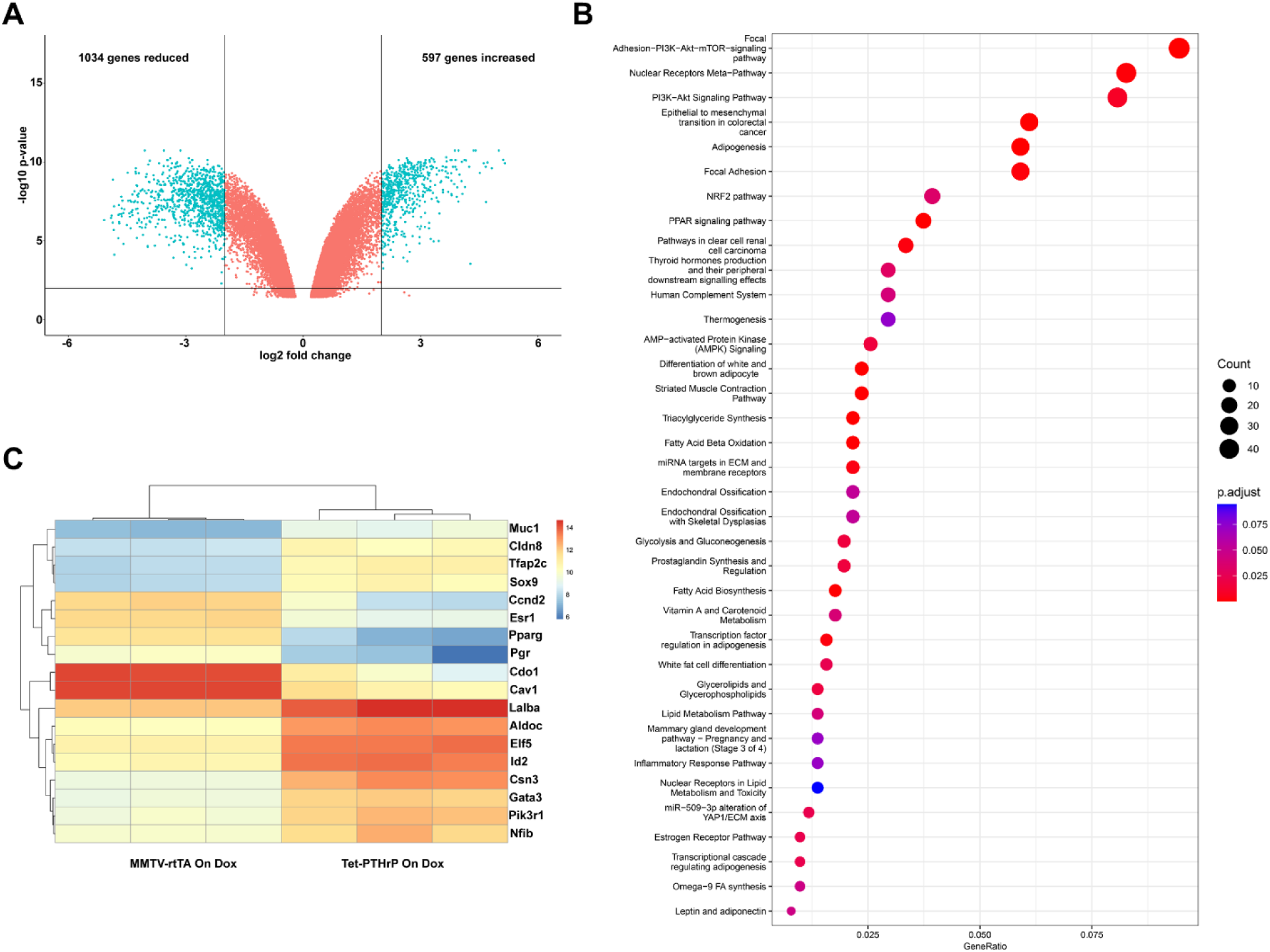
PTHrP induces changes in genes involved in secretory differentiation in luminal cells. Global mRNA profiling was performed in FACS sorted luminal mammary epithelial cells isolated from 4.5-week-old MMTV-rtTA and Tet-PTHrP mice on Dox from birth. A) Volcano plot shows the log2 fold change and variance for all transcripts in Tet-PTHrP cells relative to controls. Lines illustrate 2-fold changes and a padj of 0.01. Differentially expressed transcripts are highlighted in light blue and the number of genes increased or decreased is indicated. B) Heatmap showing relative expression change of representative genes involved in mammary gland secretory differentiation. C) Pathway analysis on differentially expressed genes. Node size represents gene count; node color represents padj.

### Overexpression of PTHrP accelerates tumor formation in MMTV-PyMT mice

We observed a cohort of 6 Tet-PTHrP mice on Dox for over a year (median 417 days) to determine whether the alveolar hyperplasia associated with PTHrP overexpression would result in the formation of mammary tumors. Only 1 mouse developed a tumor at 354 days, suggesting that PTHrP, itself, was not an efficient or dominant oncoprotein. However, in order to determine whether PTHrP overexpression might influence tumor formation caused by an established oncogene, we bred the MMTV-PyMT transgene onto Tet-PTHrP mice to generate MMTV-rtTA;TetO-PTHrP;MMTV-PyMT (Tet-PTHrP;PyMT) mice [63]. Continuous PTHrP overexpression from birth led to a dramatic acceleration of tumor formation (Fig. 4A). Microscopic tumors developed in Tet-PTHrP;PyMT mice as early as 5-10 days of age (Additional File 3A) and 100% of the mice had palpable masses in all mammary glands by just over 20 days of age (median latency of 24 days) (Fig. 4A). In contrast, control Tet-PTHrP;PyMT mice maintained off Dox developed tumors in only some glands between 40-90 days with a median latency of 71 days (Fig. 4A). PTHrP overexpression also dramatically shortened survival (Fig. 4B). When treated with Dox, Tet-PTHrP;PyMT mice became systemically ill, developed high circulating PTHrP and calcium levels (Fig. 4C), and died before 40 days of age (median survival of 30 days). In contrast, control Tet-PTHrP;PyMT mice off Dox appeared generally healthy, had normal PTHrP and calcium levels, but were euthanized due to tumor size between 50 and 100 days of age (median survival of 94 days). Importantly, overexpression of PTHrP did not increase the expression of the MMTV-PyMT transgene in cells isolated from Tet-PTHrP;PyMT tumors (Additional File 3B), demonstrating that acceleration of tumorigenesis was caused by PTHrP and not by increased PyMT expression.

**Figure 4.**
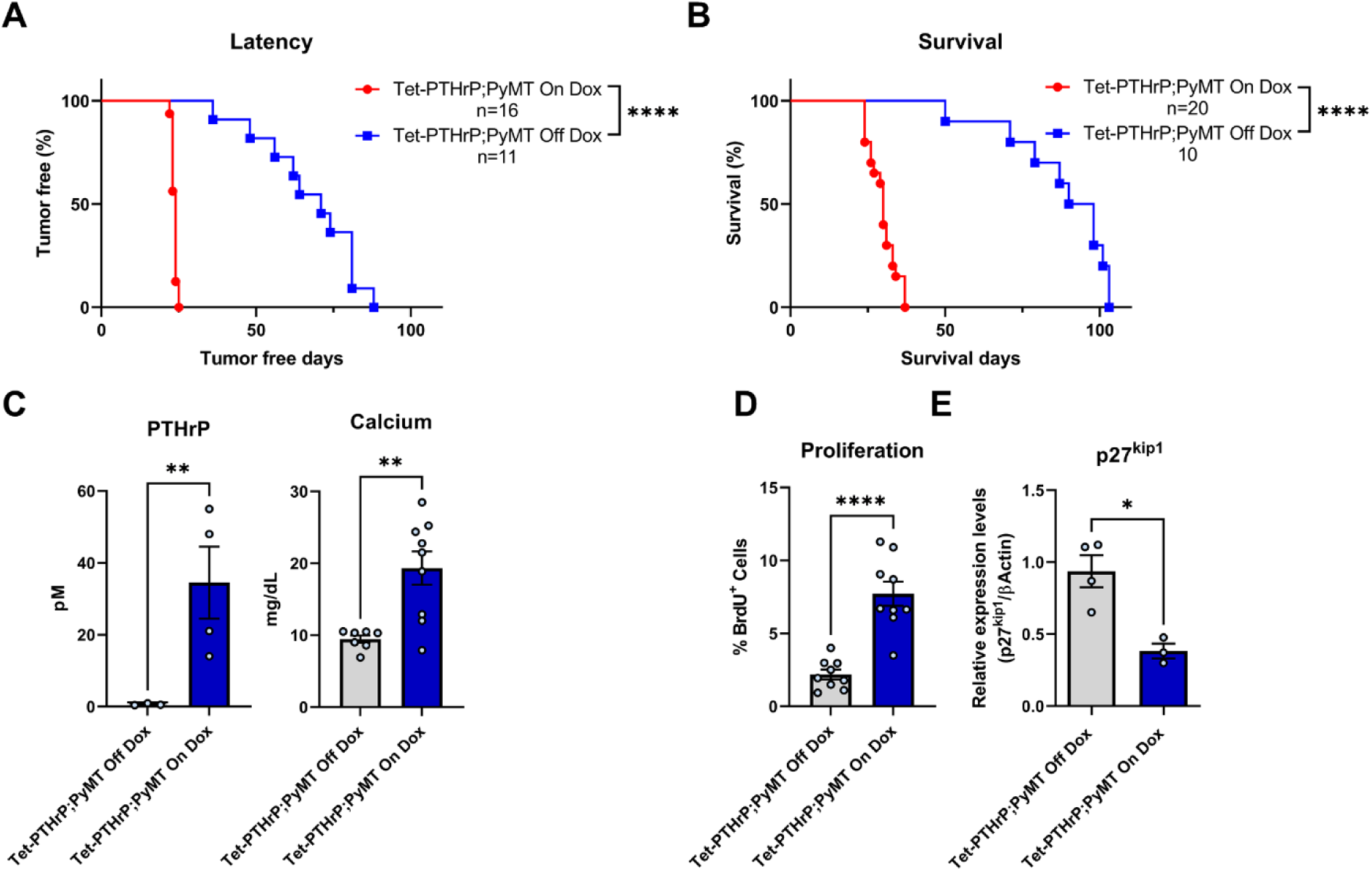
Overexpression of PTHrP accelerates tumor formation in MMTV-PyMT mice. Kaplan-Meier analysis of (A) tumor onset and (B) survival in Tet-PTHrP;PyMT mice on Dox (red) vs Tet-PTHrP;PyMT mice off Dox (blue) . (C) Circulating levels of plasma PTHrP and serum calcium concentration. (D) Quantitation of BrdU incorporation in tumor sections. Results are expressed as the percentage of BrdU-positive cells over a minimum of 1000 total cells. (E) Expression levels of *p27kip1* mRNA relative to *β-actin* mRNA in tumors. Bars represent mean ± SEM, a minimum of n=3 per group, ****p<0.0001 **p<0.01 *p<0.05.

BrdU staining demonstrated that PTHrP increased cell proliferation in mammary tumors from Tet-PTHrP;PyMT mice on Dox (Fig. 4D). Previous work has shown that PTHrP can regulate G1-S cell-cycle progression in vascular smooth muscle and breast cancer cells by modulating expression of the cell-cycle inhibitor, p27kip1 [42, 64, 65]. Therefore, we examined p27kip1 levels in tumors harvested from Tet-PTHrP;PyMT mice on or off Dox and found that increasing PTHrP production decreased p27kip1 levels in mammary tumors (Fig. 4E).

### PTHrP overexpression leads to secretory differentiation of MMTV-PyMT tumor cells

Histologically, the tumors from Tet-PTHrP;PyMT mice on Dox displayed a papillary phenotype and prominent intracellular lipid droplets as well as secretory material in extracellular “lumens” between fronds of tumor cells (Fig 5A and Additional File 4A). In addition, tumor dissection often revealed the presence of viscous white fluid resembling milk (Additional File 4B). These changes were reminiscent of the secretory differentiation seen in MECs of non-tumor bearing mice overexpressing PTHrP (Fig. 2). Therefore, we performed immunohistochemistry to examine the same mammary differentiation markers in tumor cells (Fig. 5A). Interestingly, tumors from control MMTV-PyMT mice on Dox demonstrated low levels of immunostaining for β-casein, NPT2b and pSTAT5, although expression of all three of these markers was significantly upregulated in tumors from Tet-PTHrP;PyMT mice on Dox. In addition, NKCC1 expression was downregulated by PTHrP expression. Nuclear staining for ELF5 appeared more prominent in tumors overexpressing PTHrP but nuclear staining for NF1B appeared unchanged. Western blot analyses from whole tumor lysates demonstrated similar findings. Tumors from Tet-PTHrP;PyMT mice on Dox displayed significantly higher levels of pSTAT5, β-casein and NPT2b than tumors from either MMTV-PyMT mice on Dox or from Tet-PTHrP;PyMT mice off Dox (Fig. 5B). ELF5 and NF1B levels in tumors taken from Tet-PTHrP;PyMT mice on Dox were not statistically significantly different from controls. QPCR from whole tumors revealed a significant elevation of *Wap*, *Lalba* and the different casein mRNA levels in response to PTHrP overexpression. There was also a small increase in *Elf5* gene expression but no change in *Nf1b* or *Gata3* gene expression. Overall, these changes mirrored the activation of secretory differentiation induced by PTHrP in normal MECs without the PyMT oncogene.

**Figure 5.**
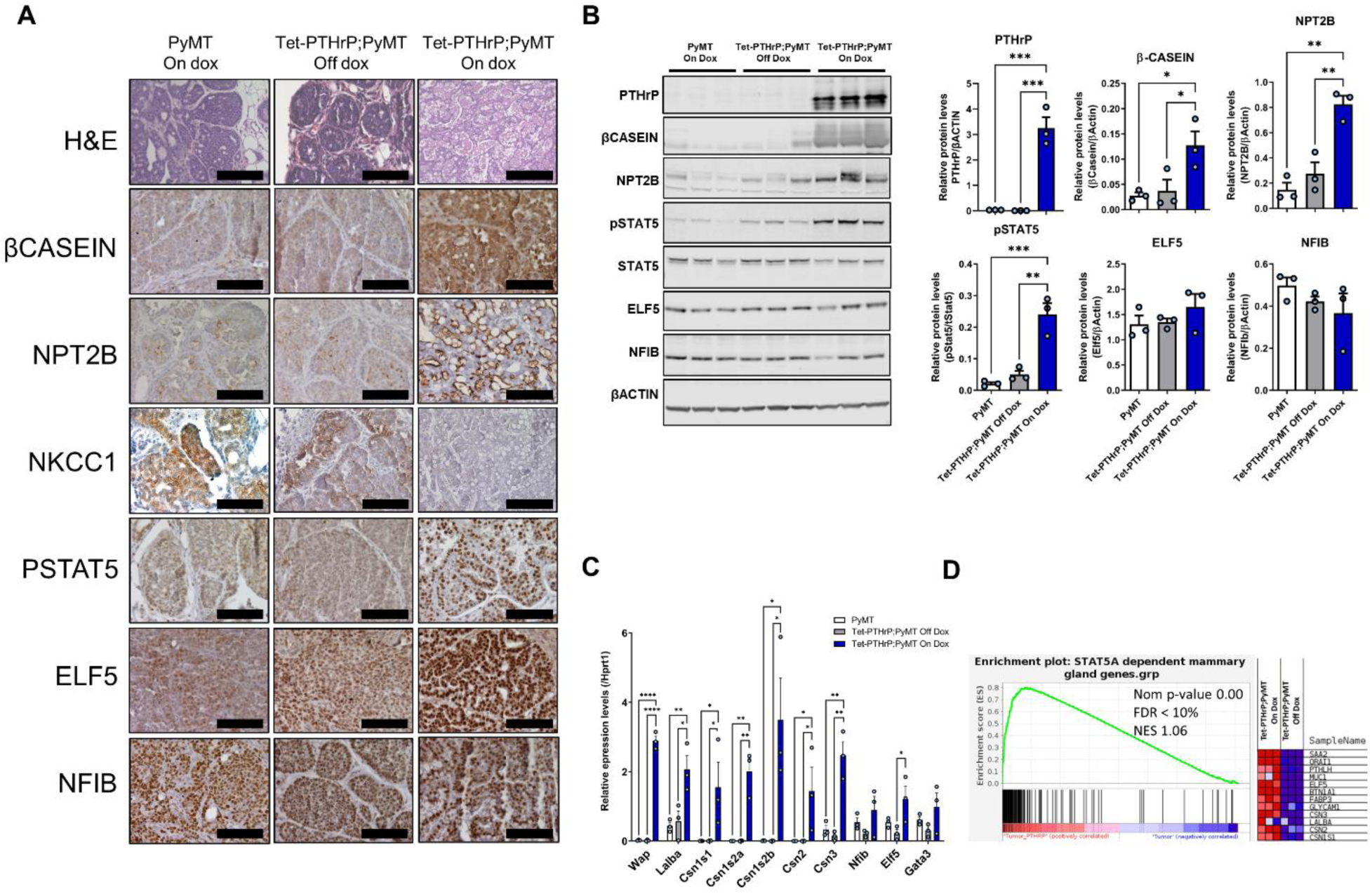
PTHrP overexpression causes secretory differentiation and Stat5 activation in PyMT tumors. (A) H&E and immunohistochemical analysis of tumors from Tet-PTHrP;PyMT mice on Dox and controls. Representative images from each group are shown. N=3. (B) Western Blots on protein lysates from whole tumors. Left shows Samples from three different mice per group were run with β-Actin as the loading control. Right shows the densitometric quantification of western blots. (C) QPCR analysis for the indicated genes of RNA isolated from whole tumors. *Hprt1* was used as housekeeping gene. N=3. D) Left shows the custom GSEA for STAT5-dependent mammary gland genes comparing Tet-PTHrP;PyMT vs PyMT mice on dox. Nom *p*-value, normalized *p*-value; FDR, false discovery rate; NES, normalized enrichment score. Right, heatmap depicting relative expression change of representative STAT5-dependent genes. Bars represent mean ± SEM, ****p<0.0001 ***p<0.001 **p<0.01 *p<0.05.

Given the apparent increased level of differentiation of the cells, we also examined whether tumors in Tet-PTHrP;PyMT mice on Dox continued to demonstrate malignant behavior. First we examined lungs from these mice for metastases (Additional File 4C) and found that 6 out of 10 Tet-PTHrP;PyMT mice treated with Dox developed lung lesions. Among those, we counted an average of 6.17 lung metastasis per mice, documenting that tumor cells retained the ability to disseminate and metastasize to distant sites. We also transplanted isolated tumor cells from Tet-PTHrP;PyMT mice into mammary fat pads of WT animals. In the presence of Dox, 75% of the 8 mice receiving these cells developed tumors that secreted PTHrP into the circulation producing significant hypercalcemia (Additional File 4D). These data demonstrate that PTHrP overexpression did not extinguish the tumor-propagating potential of the cells. Therefore, while PTHrP triggered a program of secretory differentiation in PyMT tumor cells, it did so without reversing their transformed state.

### Mammary tumors overexpressing PTHrP activate gene signatures that overlap with STAT5 Signaling and Lactation

We reasoned that the development of secretory alveolar hyperplasia in the mammary glands of the Tet-PTHrP mice and the formation of secretory adenocarcinomas in Tet-PTHrP;PyMT mice are likely to be a consequence of the activation of common STAT5-dependent pathways. To test this hypothesis, we performed a second microarray using RNA isolated from tumors of Tet-PTHrP;PyMT mice on or off Dox. As before, we used a LFC cutoff of 2 and an adjusted p value of 0.01, and identified a total of 921 differentially expressed genes (686 reduced and 235 increased) in response to PTHrP overexpression (Additional File 5A). Pathway analysis demonstrated that the differentially expressed transcripts could be grouped into pathways involved in adipocyte differentiation, fatty acid metabolism and the mammary gland transition from pregnancy to lactation, among others (Additional File 5B). We then asked specifically whether mammary epithelial cell, STAT5-dependent genes were activated in tumors overexpressing PTHrP by comparing the differentially expressed genes from PTHrP-overexpressing tumors to a previously validated set of ∼200 STAT5-dependent genes specific to mammary tissue [38]. Gene set enrichment analysis demonstrated that overexpression of PTHrP led to a significant enrichment of Stat5-dependent mRNAs in PyMT-derived tumors (Fig. 5D). To illustrate the induction of STAT5-dependent genes, the accompanying heatmap shows relative changes in the expression of 12 selected STAT5 target genes that are normally induced during lactation and are also induced by PTHrP expression in tumors. The complete list of the genes from the set and their relative change in expression in response to PTHrP overexpression is shown in Additional File 5C.

Given the similarities in the secretory phenotypes induced by PTHrP in normal MECs and in PyMT tumors, we directly compared differentially expressed genes (DEGs) in PTHrP-overexpressing tumors from Tet-PTHrP;PyMT mice with DEGs in PTHrP-overexpressing MECs from Tet-PTHrP mice (Fig. 6). There were 921 DEGs in PTHrP-overexpressing tumors and 1,631 DEGs in PTHrP overexpressing luminal MECs. Comparing these sets of genes documented a substantial overlap with a shared group of 652 genes (524 reduced and 128 increased) that were differentially expressed in both settings. Analysis of the genes in the overlap showed expression of genes involved in mammary gland development skewed toward lactation, as indicated by upregulation of *Elf5* and *Ttc* and downregulation of *Cebpα*, *Cav-1* and progesterone receptor (*Pgr*) (Fig 6B). Stimulation of ELF5 and downregulation of Cav-1, PGR, and changes in the PI3K/Akt pathway are all consistent with an increase in STAT5 signaling [35, 38, 59–62, 66, 67]. Overall, these results are consistent with the idea that PTHrP overexpression leads to STAT5 activation and secretory differentiation in both normal MECs as well as in PyMT-induced mammary tumors.

**Figure 6.**
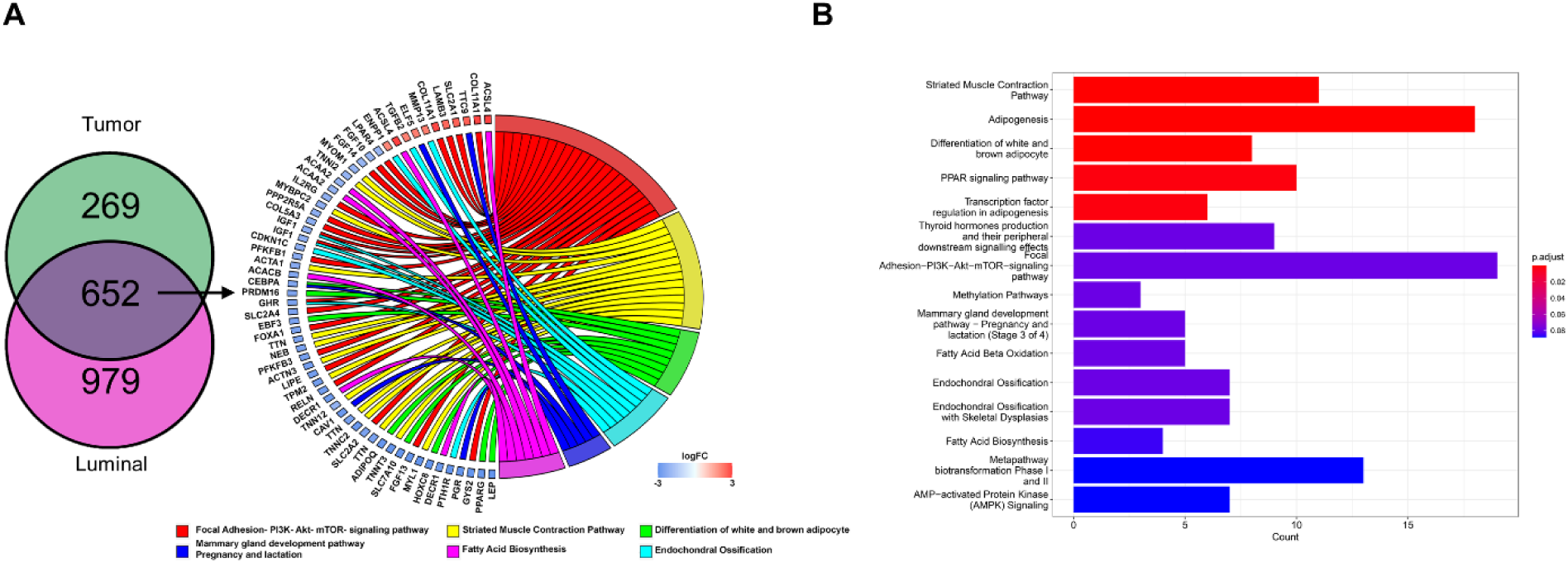
Identification of overlapping genes in mammary epithelial cells and mammary tumors overexpressing PTHrP. A) Venn diagram indicating overlap between differentially expressed genes in PTHrP-overexpressing MECs (purple) and tumors (green). B) Chord plot illustrating a detailed relationship between the log_2_-fold change (log_2_FC) of overlapped DEGs (left semicircle) and their enriched selected biological pathways. C) Pathways analysis on differentially expressed overlapped genes. Bar length represents gene count; bar color represents padj.

### Activation of Stat5 in tumor cells by PTHrP is cell autonomous and independent of PTH1R

PTHrP can signal through autocrine/paracrine mechanisms involving the activation of its cell surface receptor (PTH1R) or, alternatively, through intracrine/nuclear mechanisms [4, 5, 7]. Tran and colleagues had previously described correlations between immunostaining for nuclear PTHrP and nuclear pSTAT5 expression in human breast tumors [11]. Therefore, we hypothesized that PTHrP triggered secretory differentiation in breast cancer cells through an intracrine pathway involving Stat5 activation. In order to test this idea, we treated cells derived from mammary tumors from Tet-PTHrP;PyMT mice either with vehicle, Dox to induce endogenous PTHrP expression or with exogenous PTHrP (100nM) added to the media. Treating the cells with Dox caused increased pStat5 levels as assessed by Western analysis, whereas adding PTHrP to the media of the cells did not (Fig. 7A). In both circumstances, cells were cultured in the absence of prolactin. Similarly, Dox treatment was associated with an increase in the expression of various milk proteins by QPCR including *Csn1s1*, *Csn1s2a*, *Csn2* and *Csn3*, which, again, was not reproduced by treatment with exogenous PTHrP (Fig. 7B). Finally, we examined cell proliferation as assessed by BrdU incorporation. Induction of PTHrP with Dox treatment led to an increase in proliferation of the cells while treatment with exogenous PTHrP did not (Fig. 7C). These results suggest that the effects of PTHrP are cell autonomous, independent of prolactin stimulation, and mediated by intracrine actions of PTHrP.

**Figure 7.**
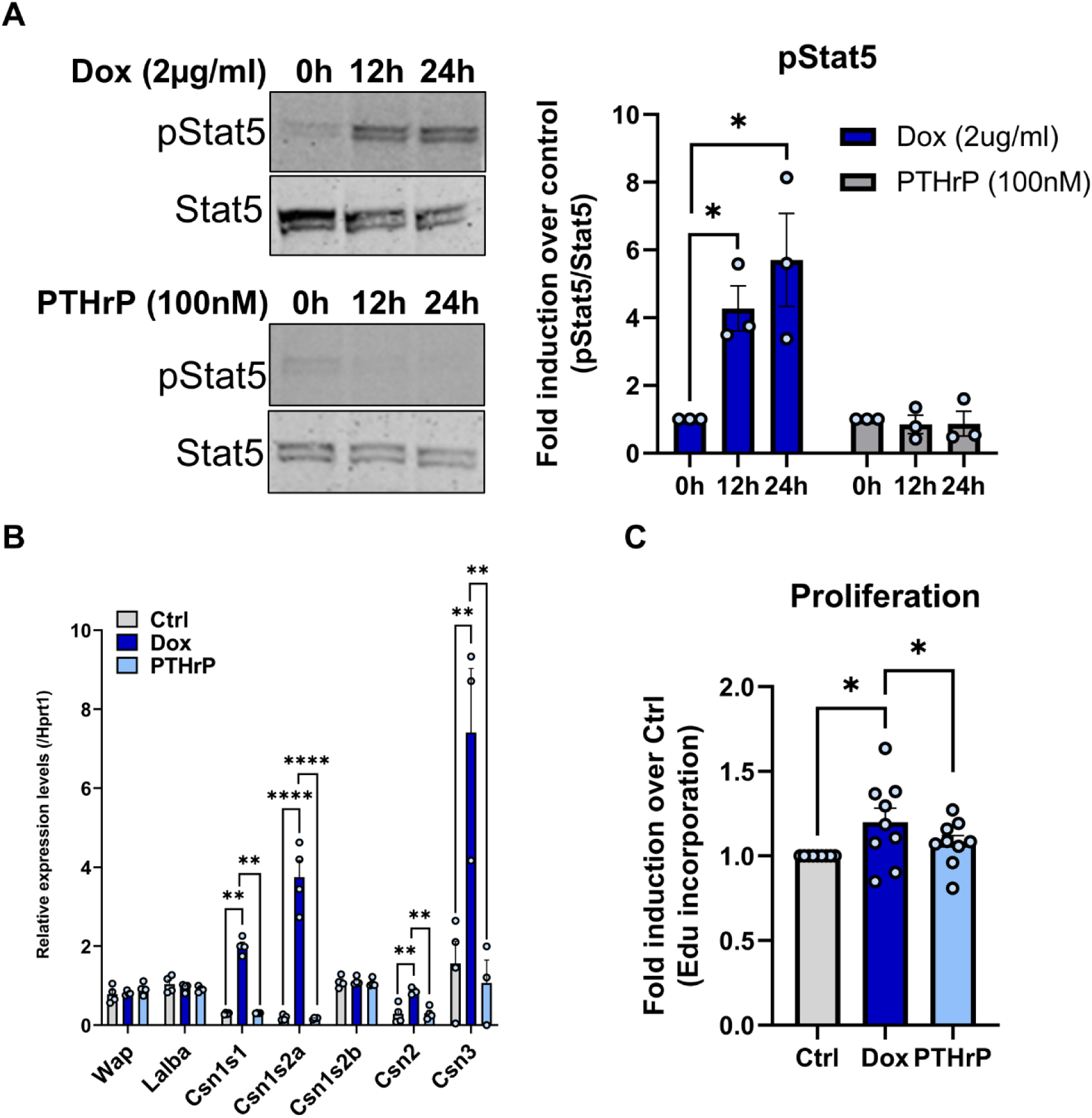
Overexpression of PTHrP, but not exogenously added PTHrP, activates Stat5 in tumor cells. Tet-PTHrP;PyMT tumor cells were treated with Dox (2µg/ml) or PTHrP 1-34 (100nM) and protein lysates and RNA was prepared. A) Western blot analysis of protein lysates. Left, representative immunoblots of p(Tyr694)Stat5 and total Stat5 are shown. Right, densitometric quantification of the western blots. N=3. B) QPCR analysis of the indicated milk proteins. *Hprt1* was used as housekeeping gene. N=3 C) Edu incorporation in cultured Tet-PTHrP;PyMT tumor cells in response to Dox or PTHrP treatment vs control. N=9. Bars represent mean ± SEM, ****p<0.0001 **p<0.01 *p<0.05.

Although the PTH1R is expressed at only very low levels, if at all, on normal mammary epithelial cells, it is expressed in many breast cancer cells [29, 68, 69]. Consistent with this literature, we found very low levels of *Pth1r* expression in non-transformed HC11 mouse mammary epithelial cells or in the mammary glands of Tet-PTHrP mice off Dox (Fig. 8A). In comparison, we found increased levels of *Pth1r* expressed in PyMT tumors from either MMTV-PyMT mice or from Tet-PTHrP;PyMT mice off Dox. Therefore, we next tested whether the effects of PTHrP on tumor cell growth and differentiation *in vivo* depended on signaling through the PTH1R by engineering mice with MMTV-Cre-mediated disruption of the *Pth1r* gene in the setting of tetracycline-regulated PTHrP overexpression and MMTV-PyMT-mediated mammary tumorigenesis. MMTV-rtTA;TetO-PTHrP; MMTV-PyMT; MMTV-Cre;PTH1R^lox/lox^ (Tet-PTHrP;PyMT;Cre;PTH1RLox) mice on Dox were followed for the development of mammary tumors and compared to Tet-PTHrP;PyMT;PTH1RLox mice that lacked the Cre transgene. As shown in Fig. 8B, the incidence and latency of tumors as well as survival in Tet-PTHrP;PyMT;Cre;PTH1RLox mice was no different than in Tet-PTHrP;PyMT;PTH1RLox mice. Tumor cells isolated from these mice demonstrated successful reduction of *Pth1r* mRNA levels (Fig. 8C), but lack of PTH1R expression had no effect on the expression of typical markers of lactation such as *Wap*, *Lalba* and multiple casein genes in whole tumor lysates (Fig 8D). In addition, ablation of the PTH1R had no effect on the expression of pSTAT5 in the nuclei of tumor cells (Fig. 8E). We further confirmed that the effects of PTHrP were independent of the PTH1R by treating Tet-PTHrP;PyMT mice with a blocking antibody against the PTH1R (anti-PTH1R) or IgG control at the same time Dox was provided (Fig. 8F-I). As expected, Tet-PTHrP;PyMT mice on Dox and treated with IgG developed hypercalcemia. However, Tet-PTHrP;PyMT mice on Dox and treated with anti-PTH1R antibody had normal calcium levels despite persistently elevated PTHrP levels, indicating that this treatment is highly effective in blocking systemic PTH1R signaling (Fig. 8F&G). In contrast, treatment with anti-PTH1R antibody did not prevent the induction of milk protein gene expression or STAT5 activation in tumor cells, demonstrating that the PTH1R is not required for PTHrP to trigger secretory differentiation in tumors (Fig. 8H&I). These results are consistent with the experiments *in vitro* suggesting that PTHrP acts in an intracrine manner.

**Figure 8.**
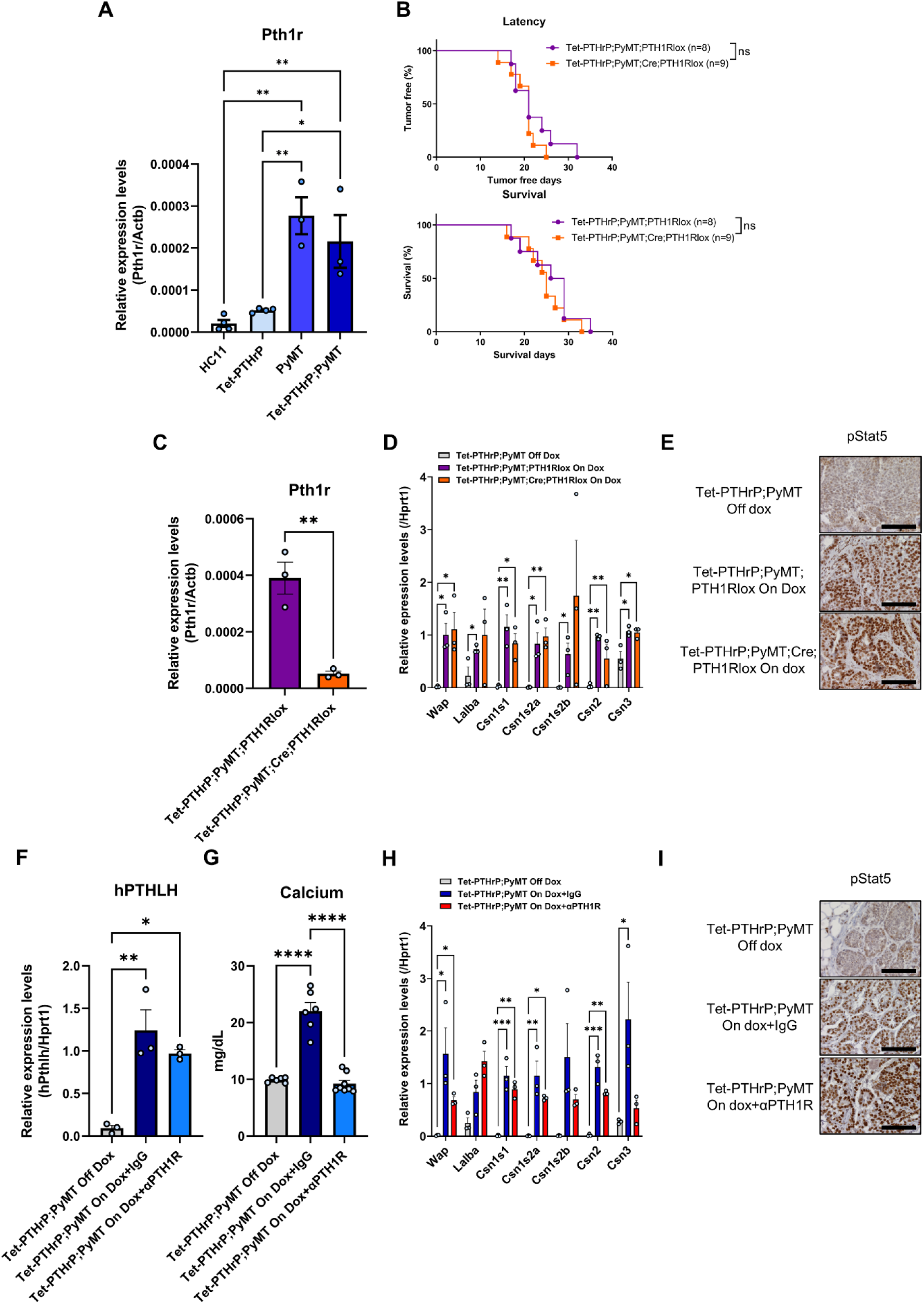
Knocking down or blocking PTH1R in tumors does not prevent the effects of PTHrP overexpression *in vivo*. A) QPCR analysis of PTH1R expression in RNA from cultured HC11, Tet-PTHrP, PyMT and Tet-PTHrP;PyMT cells. *Actb* was used as housekeeping gene. A minimum of n=3 per group. B) Kaplan-Meier analysis of tumor onset (top) and survival (bottom) of Tet-PTHrP;PyMT;PTH1RLox and Tet-PTHrP;PyMT;Cre;PTH1RLox mice treated with Dox from birth. C) Relative expression of PTH1R in RNA from isolated tumor cells. N=3. D)&H) QPCR analysis of indicated genes. *Hprt1* was used as housekeeping gene. N=3. E)&I) Representative immunohistochemical staining for nuclear pStat5 in tumor sections from Tet-PTHrP;PyMT;Cre;PTH1RLox mice treated with Dox and controls and from Tet-PTHrP;PyMT treated with Dox and an anti-PTH1R antibody (αPTH1R) and controls. N=3, Scale bar 100µm. F) QPCR analysis of hPTHLH expression in RNA from whole tumors. *Hprt1* was used as housekeeping gene. N=3. G) Serum calcium concentration. A minimum n=6. G) Bars represent mean ± SEM, ****p<0.0001 ***p<0.001 **p<0.01 *p<0.05.

### *PTHLH* Gene Expression in Human Breast Cancer Cells Correlates with Increased Expression of Genes Associated with Secretory Differentiation

In order to determine whether PTHrP production correlated with activation of STAT5-dependent, secretory differentiation pathways in human breast cancer, we examined recently published, single-cell sequencing data derived from 27 different human breast tumors (8 TNBC, 6 HER2-pos, 13 ER-pos) [70]. We were able to define *PTHLH-*high and *PTHLH-*low subsets from the pooled sequencing data of 86,277 individual epithelial tumor cells (Fig. 9A). The *PTHLH*-high cells were a distinct minority of the total cells and could be found at low levels in the three tumor sub-types. However, TNBCs had significantly more *PTHLH*-high cells (8.92%) than either HER2-positive (1.55%) or ER-positive (1.5%) tumors (Fig. 9B). We then defined the DEGs in *PTHLH-*high vs. *PTHLH*-low cells using pooled data from all tumor sub-types and performed functional pathways analyses using GSEA. We found that the DEGs were enriched in several pathways known to regulate aspects of lactation and milk production including protein secretion, fatty acid metabolism, PI3K-AKT-MTOR signaling and cholesterol homeostasis, as well as mitotic cell cycle and cell division processes (Fig. 9C). Importantly, GSEA confirmed that DEGs in *PTHLH*-high vs. *PTHLH*-low cells were significantly enriched in the hallmark IL-2-STAT5 signaling pathway [71]. Overall, these results suggest that high expression of PTHrP in human breast cancer cells is associated with expression of genes involved in milk production and STAT5 signaling.

**Figure 9.**
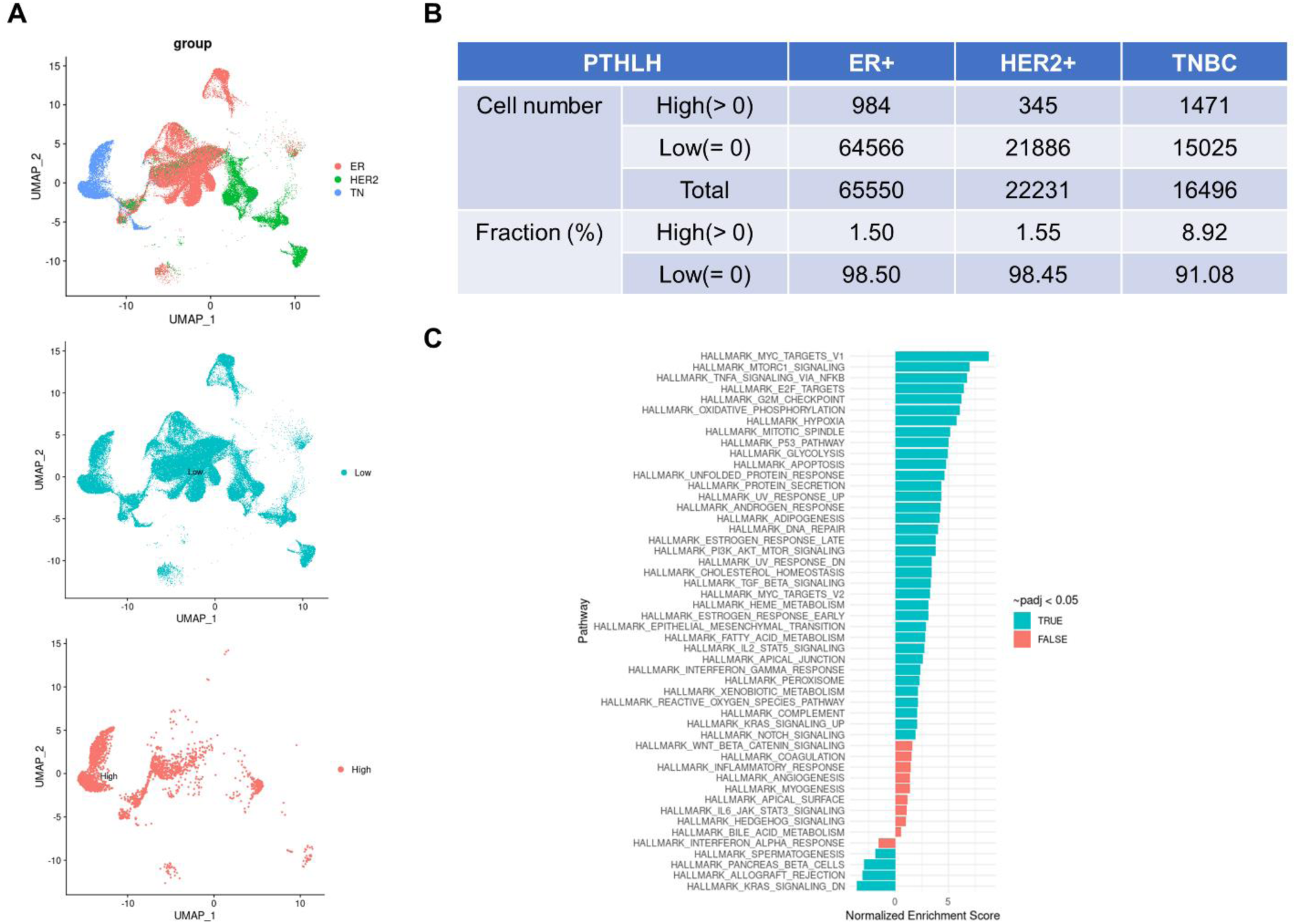
scRNA-seq analysis of PTHLH expression in human breast tumors. A) Top panel: UMAP of clusters identified by scRNAseq of epithelial cells only (EpCAM+) separated by cancer subtype (ER+ epithelial cells (n = 65,550), HER2+ epithelial cells (n = 22,231), and TNBC epithelial cells (n = 16,496)). Middle and Bottom Panel: UMAP overlays of PTHLH low and high expressing cells. B) Table containing the proportion of PTHLH expressing cells in each cancer subtype. C) fGSEA pathway analysis on DEGs from PTHLH-high vs. PTHLH-low cells using pooled data from all tumor sub-types.

## Discussion

The data presented in this study demonstrate that overexpression of PTHrP in mammary epithelial cells activates a program of secretory differentiation. When treated with Dox to induce human PTHrP (1-141) expression, the mammary glands of virgin, Tet-PTHrP mice develop alveolar hyperplasia, express histological markers of secretory differentiation, activate milk protein gene expression, and accumulate intracellular lipids. These secretory changes are accompanied by the phosphorylation of STAT5 and an increase in the expression of ELF5, two pioneering transcription factors well described to cooperate in driving gene expression necessary for milk production [35–38, 59]. Consistent with the activation of these transcription factors, we found that PTHrP upregulates patterns of gene expression previously associated with lactation. Alveolar hyperplasia and the expression of secretory differentiation markers are significantly reversed in response to the withdrawal of Dox, suggesting that they depend on the continuing presence of PTHrP. However, previous results from our lab demonstrated that, although PTHrP expression is normally activated during lactation, mammary gland specific ablation of PTHrP affects systemic calcium metabolism during lactation but does not interfere with alveolar development or with overall milk secretion [17, 43]. Given the importance of lactation to mammalian reproduction, it is not surprising that the pathways leading to secretory differentiation would be redundant. Nevertheless, these new data clearly demonstrate that PTHrP is sufficient to induce secretory differentiation in luminal epithelial cells in the absence of pregnancy.

PTHrP overexpression also drives secretory differentiation in tumor cells in the MMTV-PyMT model of breast cancer. Tet-PTHrP;PyMT mice continuously exposed to Dox develop tumors in all mammary glands by 3-4 weeks of age, a dramatic decrease in tumor latency in comparison to Tet-PTHrP;PyMT mice not treated with Dox. While PTHrP promoted premature growth of PyMT-associated mammary tumors, overexpression of PTHrP alone did not efficiently induce tumors. Therefore, in this setting, PTHrP appeared primarily to promote tumor growth rather than initiate transformation. The decrease in tumor latency was accompanied by increased rates of proliferation in the tumors. As noted previously in vascular smooth muscle cells and in human and murine breast tumor cells, increased proliferation was associated with decreased expression of the cell cycle inhibitor, p27Kip1 [31, 72]. This acceleration of tumor development is consistent with the reciprocal results of Li et al., who showed that ablation of PTHrP expression in MMTV-PyMT tumors slowed tumor growth and was associated with reduced proliferation and increased apoptosis [32]. They are also consistent with prior results from our group demonstrating that ablation of the CaSR in MMTV-PyMT tumors or in human BT474 and MDA.MB-231.1833 cells reduced PTHrP expression, which was associated with a reduction in proliferation and increased expression of p27kip1 [31]. Thus, although there have been variable reports on the effects of PTHrP on human breast cancer cell lines, in mice, PTHrP clearly promotes proliferation in mammary tumor cells expressing PyMT.

While PTHrP accelerates the growth of PyMT tumors, it also promotes secretory differentiation. This is associated with an increase in the expression of differentiation markers, milk protein genes, Elf5, and pSTAT5. Despite an apparent shift to a more differentiated state, tumors continued to metastasize, and cells derived from the tumors overexpressing PTHrP were able to form new tumors when transplanted into non-transgenic mice. The histological appearance of the tumors in Tet-PTHrP;PyMT mice, their expression of pSTAT5 and the accumulation of milk-like secretions is reminiscent of a rare type of human breast cancer known as “secretory carcinoma of the breast” [73–77]. The majority of these tumors have been shown to contain t(12;15)(p13;q25) chromosomal translocations that results in a fusion oncogene (ETV6-NTRK3) consisting of the oligomerization domain of ETV6 fused to the protein tyrosine kinase domain of the neurotropin 3 receptor (NTRK3). Although most secretory carcinomas behave in an indolent manner, some patients developed metastatic lesions. When an ETV6-NTRK3 construct was knocked into mice, they developed mammary alveolar hyperplasia followed by the development of multifocal tumors with short latency, again reminiscent of the effects of overexpressing PTHrP on PyMT-mediated tumorigenesis [76]. Although there is no known link between PTHrP expression and the expression or activity of NTRK3 or other neurotropin receptors, it has been suggested that the transforming ability of the ETV6-NTRK3 fusion oncogene relies on activation of the AP1 transcription complex [76]. Given the similarities between PTHrP overexpression in PyMT tumors and this model of secretory carcinomas, as well as the fact that PTHrP has been shown to activate AP1 signaling by increasing c-fos and/or JunB expression in several cell types, further study of potential interactions between PTHrP and AP1 signaling in breast cancer may be revealing [78, 79].

Multiple lines of evidence suggest that the effects of PTHrP on activating secretory differentiation pathways as well as on promoting tumor cell proliferation are mediated by an intracrine pathway rather than through its cell surface receptor. First, previous experiments overexpressing PTHrP in mammary gland myoepithelial cells did not lead to alveolar hyperplasia and secretory differentiation although, similar to the results reported here, it did inhibit ductal elongation during puberty [56, 68]. These differences are not compatible with a typical paracrine mode of action given that the 2 cell types overexpressing PTHrP in these different models are adjacent to each other. Instead, the different phenotypes in these models suggest a cell-autonomous and cell-restricted mechanism of action drives the secretory differentiation. Second, in cells derived from Tet-PTHrP;PyMT mammary tumors, inducing PTHrP expression by treating them with Dox stimulates cell proliferation, activates STAT5 and increases milk proteins gene expression, but treating the cells with exogenous PTHrP does not. Thus, PTHrP is sufficient to induce secretory differentiation, but only if produced within the tumor cells, suggesting a cell autonomous and intracrine mechanism. Third, reducing PTH1R expression in tumor cells does not alter tumor growth or secretory differentiation of the tumor cells, demonstrating that tumor expression of the PTH1R is not required for the observed phenotype. Lastly, treating tumor-bearing Tet-PTHrP;PyMT mice with anti-PTH1R antibodies corrects hypercalcemia but does not reverse STAT5 activation or reduce the expression of secretory markers, demonstrating that secreted PTHrP does not act systemically or on non-tumor cells in the microenvironment to induce paracrine cascades supporting secretory differentiation. These results are consistent with the observations of Tran et al., who previously reported that nuclear PTHrP staining correlates with nuclear pSTAT5 staining in human breast cancers [11]. In addition, Johnson et al. showed that PTHrP overexpression in MCF7 cells results in the downregulation of several pro-dormancy genes and suggested that these actions may occur through PTH1R-independent actions [69]. Finally, prior results from our laboratory have demonstrated that intracrine/nuclear actions of PTHrP downstream of the calcium-sensing receptor are important in modulating cell proliferation and survival in human breast cancer cell lines and in PyMT-induced mouse mammary tumors [31].

PTHrP is widely recognized to be important for the progression of osteolytic bone metastases from breast cancer [21, 80], although its role in the initiation, growth or progression of primary breast tumors is less clear. The results we report here agree with those of Li et al., demonstrating that PTHrP stimulates mammary tumor progression and results in shorter survival in MMTV-PyMT mice [32]. As compared to studies in mice, PTHrP has been variably suggested to either promote or to inhibit breast cancer cell proliferation, differentiation and death in human breast cancer cell lines [5, 6, 11, 31, 34]. Likewise, studies examining PTHrP staining in human breast cancers have reported differing correlations between PTHrP and tumor behavior. Some studies have reported that PTHrP expression correlates with estrogen receptor and progesterone receptor expression, a more differentiated histology, fewer metastases and a better prognosis [11, 27]. In contrast, other studies have suggested that increased PTHrP expression predicts worse survival and increases brain or bone metastases when measured in all breast tumors, in triple-negative breast cancers or in circulating tumor cells [26, 29, 81, 82]. One possible explanation for these conflicting results may be related to differing effects of PTHrP in luminal vs. triple negative sub-types of breast cancer. Another may relate to our observation that PTHrP overexpression results in the upregulation of STAT5 activation. STAT5 is critical to the proliferation and secretory differentiation of normal breast epithelial cells during pregnancy and lactation, but it seems to mirror PTHrP in having different effects on tumor progression in mice and humans. Loss of STAT5 impedes the development of tumors in T-antigen-dependent mouse models, while overexpression of wild-type or constitutively active STAT5 accelerates tumor formation in these models [37, 83–85]. By contrast, the activation of STAT5 in human breast cancers has generally been observed to be an indicator of more differentiated tumors and a better prognosis [11, 83, 86]. Our findings and those of Tran et al mirror the previous literature in that PTHrP expression increases STAT5 and tumor progression in MMTV-PyMT mice, but PTHrP expression correlates with nuclear STAT5 expression and better outcome in human breast cancer. This may not be the entire bottom line given the recent report from Assaker and colleagues suggesting that tumor PTHrP expression at the time of diagnosis correlated with subsequent brain metastases and poor survival in patients with triple negative breast cancer (TNBC) [81]. Interestingly, we found the highest numbers of cells with elevated PTHrP gene expression in TNBC’s using single cell sequencing data. Furthermore, genes potentially involved in Stat5 signaling were enriched in TNBC cells expressing higher levels of PTHrP. Therefore, it is possible that interactions between PTHrP and Stat5 may have different consequences depending on the sub-type of breast cancer. Sorting out the details of when and how PTHrP affects different breast cancers in different fashions will be critical to understanding the reported association between the *PTHLH* gene and breast cancer in GWAS studies [23–25].

## Conclusions

In summary, we report that PTHrP overexpression activates STAT5, increases Elf5 expression, and leads to increased proliferation and secretory differentiation of both normal luminal mammary epithelial and mammary tumor cells in mice. This is the result of an intracrine pathway rather than a function of secreted PTHrP. We also find the greatest proportion of individual human tumor cells with high PTHrP expression in triple negative breast cancers, where higher PTHrP levels correlate with an enrichment of STAT5-related gene expression. Further work to better understand how intracrine PTHrP signaling interacts with STAT5 signaling may help to resolve conflicting published data regarding the overall effects of PTHrP on tumor behavior and patient survival.

## Declarations

### Ethics approval and consent to participate

Not applicable

### Consent for publication

Not applicable

### Availability of data and materials

The dataset supporting the conclusions of this article is available in the ArrayExpress database (http://www.ebi.ac.uk/arrayexpress) under accession number E-MTAB-11281.

### Author contributions

DYG, KBG, FMT, LC, JJW conceived and designed research. DYG, KBG, FMT, PD, JRH, CM performed experiments. MGS, JL, JC curated and analyzed data. DYG, KBG, FMT, LC, JJW interpreted results of experiments. DYG and JJW drafted manuscript. All authors read and approved the final manuscript.

## Acknowledgments

We are grateful to Dr. Henry Kronenberg for providing PTH1R^fl/fl^ mice for our studies, to Dr. James Turner for the anti-NKCC1 antibody and to Dr. Jürg Biber for the anti-NPT2b antibody.

## List of abbreviations

PTHrP: Parathyroid hormone-related protein
PTH: Parathyroid hormone
PTH1R: PTH/PTHrP receptor 1
MMTV: mouse mammary tumor virus long terminal repeat
ELF5: E74-like factor-5
TEB: Terminal end bud
MEC: Mammary epithelial cell
NKCC1: Sodium-potassium-chloride co-transporter 1
NPT2b: Sodium-phosphate transporter 2b
WAP: Whey acidic protein
LALBA: Alpha lactalbumin
DOX: Doxycycline
NF1B: Nuclear factor 1B
GSEA: Gene set enrichment analysis
DEG: Differentially expressed gene
BrdU: Bromodeoxyuridine
EdU: 5-ethynyl-2’-deoxyuridine
TNBC: Triple negative breast cancer
ER: Estrogen receptor

## Additional material

**Additional File 1**

.tiff

**Overexpression of PTHrP causes milk production in mammary glands from virgin mice.** A) Picture of the number 4 inguinal mammary gland from virgin, 13-week-old Tet-PTHrP mouse on dox showing milk accumulation. B) Densitometric quantification of the western blots for the indicated milk proteins and secretory differentiation markers shown in Figure 5. Bars represent mean ± SEM, n=3 per group, ****p<0.0001 ***p<0.001 **p<0.01 *p<0.05.

**Additional file 2.**

.tiff

**Alveolar hyperplasia and the mature secretory phenotype require ongoing exposure to PTHrP.** Immunohistochemical staining of mammary gland sections. Representative images of an n=3 per group are shown. Scale bar 100µm.

**Additional file 3.**

.tiff

**PTHrP overexpression causes microscopic tumors in Tet-PTHrP;PyMT mice as early as 10 days of age.** A) Whole-mount analysis of carmine-stained, inguinal mammary glands from 10 day-old, Tet-PTHrP;PyMT mice on dox. Representative images of two out of three mice. Scale bars 1mm (left), 100µm (right). B) QPCR analysis of *Pymt* mRNA expression in RNA from isolated tumor cells. *Actb* was used as a housekeeping gene. Bars represent mean ± SEM, n=6, ns: not significant.

**Additional file 4.**

.tiff

**PTHrP induces secretory differentiation in PyMT tumor cells, without reversing their transformed state**. A) H&E staining of tumors from different mouse genotypes and Dox treatments as detailed on the left. Representative images from 3 different tumors and mice per group. Scale bar 100µm. B) Picture of the third and fourth mammary gland containing tumors from Tet-PTHrP;PyMT mouse on Dox showing milk accumulation. C) H&E staining of lung sections from Tet-PTHrP;PyMT mice on Dox. Black boxes highlight lung metastases. Representative images of metastasis from 3 different mice. Scale bar 100µm. D) Plasma PTHrP and serum calcium concentration from WT mice on Dox transplanted with isolated Tet-PTHrP;PyMT tumor cells. Bars represent mean ± SEM, n=6.

**Additional file 5.**

.tiff

**Global mRNA profiling in tumors of Tet-PTHrP;PyMT and PyMT mice on Dox.** A) Volcano plot shows the log_2_ fold change and variance for all transcripts in PTHrP-overexpressing tumors relative to controls. Lines illustrate 2-fold changes and a padj of 0.01. Differentially expressed transcripts are highlighted in light blue and the number of genes increased or decreased is indicated. B) Pathway analysis on differentially expressed genes. Node size represents gene count; node color represents padj. C) Heatmap of STAT5-dependent mammary gland genes comparing Tet-PTHrP;PyMT vs PyMT mice on Dox using GSEA. N=3.

## References

1. Stewart AF, Horst RL, Deftos LJ, Cadman EC, Lang R, Broadus AE: Biochemical evaluation of patients with cancer-associated hypercalcemia: evidence for humoral and nonhumoral groups. New England Journal of Medicine 1980, 303:1377–1383.

2. Strewler GJ: The physiology of parathyroid hormone-related protein. N Engl J Med 2000, 342(3):177–185.

3. Wysolmerski JJ: Parathyroid hormone-related protein: an update. J Clin Endocrinol Metab 2012, 97(9):2947–2956.

4. McCauley LK, Martin TJ: Twenty-five years of PTHrP progress: from cancer hormone to multifunctional cytokine. J Bone Miner Res 2012, 27(6):1231–1239.

5. Wysolmerski JJ: Parathyroid hormone-related protein: an update. J Clin Endocrinol Metab 2012, 97(9):2947–2956.

6. Zhang R, Li J, Assaker G, Camirand A, Sabri S, Karaplis AC, Kremer R: Parathyroid Hormone-Related Protein (PTHrP): An Emerging Target in Cancer Progression and Metastasis. Adv Exp Med Biol 2019, 1164:161–178.

7. Fiaschi-Taesch NM, Stewart AF: Minireview: parathyroid hormone-related protein as an intracrine factor--trafficking mechanisms and functional consequences. Endocrinology 2003, 144(2):407–411.

8. Jans DA, Thomas RJ, Gillespie MT: Parathyroid hormone-related protein (PTHrP): a nucleocytoplasmic shuttling protein with distinct paracrine and intracrine roles. Vitam Horm 2003, 66:345–384.

9. Miao D, Su H, He B, Gao J, Xia Q, Zhu M, Gu Z, Goltzman D, Karaplis AC: Severe growth retardation and early lethality in mice lacking the nuclear localization sequence and C-terminus of PTH-related protein. Proc Natl Acad Sci U S A 2008, 105(51):20309–20314.

10. Toribio RE, Brown HA, Novince CM, Marlow B, Hernon K, Lanigan LG, Hildreth BE, 3rd, Werbeck JL, Shu ST, Lorch G, et al. The midregion, nuclear localization sequence, and C terminus of PTHrP regulate skeletal development, hematopoiesis, and survival in mice. FASEB J 2010, 24(6):1947–1957.

11. Tran TH, Utama FE, Sato T, Peck AR, Langenheim JF, Udhane SS, Sun Y, Liu C, Girondo MA, Kovatich AJ, et al. Loss of Nuclear Localized Parathyroid Hormone-Related Protein in Primary Breast Cancer Predicts Poor Clinical Outcome and Correlates with Suppressed Stat5 Signaling. Clin Cancer Res 2018, 24(24):6355–6366.

12. Cormier S, Delezoide AL, Silve C: Expression patterns of parathyroid hormone-related peptide (PTHrP) and parathyroid hormone receptor type 1 (PTHR1) during human development are suggestive of roles specific for each gene that are not mediated through the PTHrP/PTHR1 paracrine signaling pathway. Gene Expr Patterns 2003, 3(1):59–63.

13. Wysolmerski JJ, Cormier S, Philbrick WM, Dann P, Zhang JP, Roume J, Delezoide AL, Silve C: Absence of functional type 1 parathyroid hormone (PTH)/PTH-related protein receptors in humans is associated with abnormal breast development and tooth impaction. J Clin Endocrinol Metab 2001, 86(4):1788–1794.

14. Wysolmerski JJ, Philbrick WM, Dunbar ME, Lanske B, Kronenberg H, Broadus AE: Rescue of the parathyroid hormone-related protein knockout mouse demonstrates that parathyroid hormone-related protein is essential for mammary gland development. Development 1998, 125(7):1285–1294.

15. Hiremath M, Wysolmerski J: Parathyroid hormone-related protein specifies the mammary mesenchyme and regulates embryonic mammary development. Journal of mammary gland biology and neoplasia 2013, 18(2):171–177.

16. Grinman D, Athonvarungkul D, Wysolmerski J, Jeong J: Calcium Metabolism and Breast Cancer: Echoes of Lactation? Curr Opin Endocr Metab Res 2020, 15:63–70.

17. VanHouten JN, Dann P, Stewart AF, Watson CJ, Pollak M, Karaplis AC, Wysolmerski JJ: Mammary-specific deletion of parathyroid hormone-related protein preserves bone mass during lactation. J Clin Invest 2003, 112(9):1429–1436.

18. VanHouten JN, Wysolmerski JJ: Low estrogen and high parathyroid hormone-related peptide levels contribute to accelerated bone resorption and bone loss in lactating mice. Endocrinology 2003, 144(12):5521–5529.

19. Mamillapalli R, VanHouten J, Dann P, Bikle D, Chang W, Brown E, Wysolmerski J: Mammary-specific ablation of the calcium-sensing receptor during lactation alters maternal calcium metabolism, milk calcium transport, and neonatal calcium accrual. Endocrinology 2013, 154(9):3031–3042.

20. Kim W, Wysolmerski JJ: Calcium-Sensing Receptor in Breast Physiology and Cancer. Front Physiol 2016, 7:440.

21. Akhtari M, Mansuri J, Newman KA, Guise TM, Seth P: Biology of breast cancer bone metastasis. Cancer Biol Ther 2008, 7(1):3–9.

22. Martin TJ, Johnson RW: Multiple actions of parathyroid hormone-related protein in breast cancer bone metastasis. Br J Pharmacol 2021, 178(9):1923–1935.

23. Purrington KS, Slager S, Eccles D, Yannoukakos D, Fasching PA, Miron P, Carpenter J, Chang-Claude J, Martin NG, Montgomery GW, et al. Genome-wide association study identifies 25 known breast cancer susceptibility loci as risk factors for triple-negative breast cancer. Carcinogenesis 2014, 35(5):1012–1019.

24. Teraoka SN, Bernstein JL, Reiner AS, Haile RW, Bernstein L, Lynch CF, Malone KE, Stovall M, Capanu M, Liang X, et al. Single nucleotide polymorphisms associated with risk for contralateral breast cancer in the Women’s Environment, Cancer, and Radiation Epidemiology (WECARE) Study. Breast Cancer Res 2011, 13(6):R114.

25. Zeng C, Guo X, Long J, Kuchenbaecker KB, Droit A, Michailidou K, Ghoussaini M, Kar S, Freeman A, Hopper JL, et al. Identification of independent association signals and putative functional variants for breast cancer risk through fine-scale mapping of the 12p11 locus. Breast Cancer Res 2016, 18(1):64.

26. Bundred NJ, Walls J, Ratcliffe WA: Parathyroid hormone-related protein, bone metastases and hypercalcaemia of malignancy. Ann R Coll Surg Engl 1996, 78(4):354–358.

27. Henderson MA, Danks JA, Slavin JL, Byrnes GB, Choong PF, Spillane JB, Hopper JL, Martin TJ: Parathyroid hormone-related protein localization in breast cancers predict improved prognosis. Cancer Res 2006, 66(4):2250–2256.

28. Hoey RP, Sanderson C, Iddon J, Brady G, Bundred NJ, Anderson NG: The parathyroid hormone-related protein receptor is expressed in breast cancer bone metastases and promotes autocrine proliferation in breast carcinoma cells. Br J Cancer 2003, 88(4):567–573.

29. Linforth R, Anderson N, Hoey R, Nolan T, Downey S, Brady G, Ashcroft L, Bundred N: Coexpression of parathyroid hormone related protein and its receptor in early breast cancer predicts poor patient survival. Clin Cancer Res 2002, 8(10):3172–3177.

30. Takagaki K, Takashima T, Onoda N, Tezuka K, Noda E, Kawajiri H, Ishikawa T, Hirakawa K: Parathyroid hormone-related protein expression, in combination with nodal status, predicts bone metastasis and prognosis of breast cancer patients. Exp Ther Med 2012, 3(6):963–968.

31. Kim W, Takyar FM, Swan K, Jeong J, VanHouten J, Sullivan C, Dann P, Yu H, Fiaschi-Taesch N, Chang W, et al. Calcium-Sensing Receptor Promotes Breast Cancer by Stimulating Intracrine Actions of Parathyroid Hormone-Related Protein. Cancer Res 2016, 76(18):5348–5360.

32. Li J, Karaplis AC, Huang DC, Siegel PM, Camirand A, Yang XF, Muller WJ, Kremer R: PTHrP drives breast tumor initiation, progression, and metastasis in mice and is a potential therapy target. J Clin Invest 2011, 121(12):4655–4669.

33. Luparello C: Parathyroid Hormone-Related Protein (PTHrP): A Key Regulator of Life/Death Decisions by Tumor Cells with Potential Clinical Applications. Cancers (Basel) 2011, 3(1):396–407.

34. Luparello C, Sirchia R, Lo Sasso B: Midregion PTHrP regulates Rip1 and caspase expression in MDA-MB231 breast cancer cells. Breast Cancer Res Treat 2008, 111(3):461–474.

35. Choi YS, Chakrabarti R, Escamilla-Hernandez R, Sinha S: Elf5 conditional knockout mice reveal its role as a master regulator in mammary alveolar development: failure of Stat5 activation and functional differentiation in the absence of Elf5. Dev Biol 2009, 329(2):227–241.

36. Frend HT, Watson CJ: Elf5 - breast cancer’s little helper. Breast Cancer Res 2013, 15(2):307.

37. Furth PA, Nakles RE, Millman S, Diaz-Cruz ES, Cabrera MC: Signal transducer and activator of transcription 5 as a key signaling pathway in normal mammary gland developmental biology and breast cancer. Breast Cancer Res 2011, 13(5):220.

38. Metser G, Shin HY, Wang C, Yoo KH, Oh S, Villarino AV, O’Shea JJ, Kang K, Hennighausen L: An autoregulatory enhancer controls mammary-specific STAT5 functions. Nucleic Acids Res 2016, 44(3):1052–1063.

39. Gunther EJ, Belka GK, Wertheim GBW, Wang J, Hartman JL, Boxer RB, Chodosh LA: A novel doxycycline-inducible system for the transgenic analysis of mammary gland biology. The FASEB Journal 2002, 16(3):283–292.

40. Dunbar M, Dann P, Brown C, Van Houton J, Dreyer B, Philbrick W, Wysolmerski J: Temporally regulated overexpression of parathyroid hormone-related protein in the mammary gland reveals distinct fetal and pubertal phenotypes. Journal of Endocrinology 2001, 171(3):403–416.

41. Kobayashi T, Chung U-I, Schipani E, Starbuck M, Karsenty G, Katagiri T, Goad DL, Lanske B, Kronenberg HM: PTHrP and Indian hedgehog control differentiation of growth plate chondrocytes at multiple steps. Development 2002, 129(12):2977–2986.

42. Kim W, Takyar FM, Swan K, Jeong J, Vanhouten J, Sullivan C, Dann P, Yu H, Fiaschi-Taesch N, Chang W, et al. Calcium-Sensing Receptor Promotes Breast Cancer by Stimulating Intracrine Actions of Parathyroid Hormone–Related Protein. Cancer Research 2016, 76(18):5348–5360.

43. Boras-Granic K, VanHouten J, Hiremath M, Wysolmerski J: Parathyroid hormone-related protein is not required for normal ductal or alveolar development in the post-natal mammary gland. PLoS One 2011, 6(11):e27278.

44. Kocatürk B, Versteeg HH: Orthotopic Injection of Breast Cancer Cells into the Mammary Fat Pad of Mice to Study Tumor Growth. Journal of Visualized Experiments 2015(96).

45. Yamaji D, Na R, Feuermann Y, Pechhold S, Chen W, Robinson GW, Hennighausen L: Development of mammary luminal progenitor cells is controlled by the transcription factor STAT5A. Genes & Development 2009, 23(20):2382–2387.

46. Huber W, Carey VJ, Gentleman R, Anders S, Carlson M, Carvalho BS, Bravo HC, Davis S, Gatto L, Girke T, et al. Orchestrating high-throughput genomic analysis with Bioconductor. Nature Methods 2015, 12(2):115–121.

47. Martens M, Ammar A, Riutta A, Waagmeester A, Denise, Hanspers K, Ryan, Digles D, Elisson, Ehrhart F, et al. WikiPathways: connecting communities. Nucleic Acids Research 2021, 49(D1):D613–D621.

48. Walter W, Sánchez-Cabo F, Ricote M: GOplot: an R package for visually combining expression data with functional analysis: Fig. 1. Bioinformatics 2015, 31(17):2912–2914.

49. Subramanian A, Tamayo P, Mootha VK, Mukherjee S, Ebert BL, Gillette MA, Paulovich A, Pomeroy SL, Golub TR, Lander ES, et al. Gene set enrichment analysis: A knowledge-based approach for interpreting genome-wide expression profiles. Proceedings of the National Academy of Sciences 2005, 102(43):15545–15550.

50. H W: ggplot2: Elegant Graphics for Data Analysis: Springer-Verlag; 2016.

51. Hao Y, Hao S, Andersen-Nissen E, Mauck WM, Zheng S, Butler A, Lee MJ, Wilk AJ, Darby C, Zager M, et al. Integrated analysis of multimodal single-cell data. Cell 2021, 184(13):3573–3587.e3529.

52. Pal B, Chen Y, Vaillant F, Capaldo BD, Joyce R, Song X, Bryant VL, Penington JS, Di Stefano L, Tubau Ribera N, et al. A single-cell RNA expression atlas of normal, preneoplastic and tumorigenic states in the human breast. The EMBO Journal 2021, 40(11).

53. Korotkevich G, Sukhov V, Budin N, Shpak B, Artyomov MN, Sergushichev A: Fast gene set enrichment analysis. In.: Cold Spring Harbor Laboratory; 2016.

54. Durinck S, Spellman PT, Birney E, Huber W: Mapping identifiers for the integration of genomic datasets with the R/Bioconductor package biomaRt. Nature Protocols 2009, 4(8):1184–1191.

55. Ecker BL, Lee JY, Sterner CJ, Solomon AC, Pant DK, Shen F, Peraza J, Vaught L, Mahendra S, Belka GK, et al. Impact of obesity on breast cancer recurrence and minimal residual disease. Breast Cancer Res 2019, 21(1):41.

56. Dunbar ME, Dann P, Brown CW, Van Houton J, Dreyer B, Philbrick WP, Wysolmerski JJ: Temporally regulated overexpression of parathyroid hormone-related protein in the mammary gland reveals distinct fetal and pubertal phenotypes. J Endocrinol 2001, 171(3):403–416.

57. Anderson SM, Rudolph MC, McManaman JL, Neville MC: Key stages in mammary gland development. Secretory activation in the mammary gland: it’s not just about milk protein synthesis! Breast Cancer Res 2007, 9(1):204.

58. Mukhopadhyay SS, Wyszomierski SL, Gronostajski RM, Rosen JM: Differential interactions of specific nuclear factor I isoforms with the glucocorticoid receptor and STAT5 in the cooperative regulation of WAP gene transcription. Mol Cell Biol 2001, 21(20):6859–6869.

59. Shin HY, Willi M, HyunYoo K, Zeng X, Wang C, Metser G, Hennighausen L: Hierarchy within the mammary STAT5-driven Wap super-enhancer. Nat Genet 2016, 48(8):904–911.

60. Andrechek ER, Mori S, Rempel RE, Chang JT, Nevins JR: Patterns of cell signaling pathway activation that characterize mammary development. Development 2008, 135(14):2403–2413.

61. Clarkson RW, Wayland MT, Lee J, Freeman T, Watson CJ: Gene expression profiling of mammary gland development reveals putative roles for death receptors and immune mediators in post-lactational regression. Breast Cancer Res 2004, 6(2):R92–109.

62. Rudolph MC, McManaman JL, Hunter L, Phang T, Neville MC: Functional development of the mammary gland: use of expression profiling and trajectory clustering to reveal changes in gene expression during pregnancy, lactation, and involution. J Mammary Gland Biol Neoplasia 2003, 8(3):287–307.

63. Attalla S, Taifour T, Bui T, Muller W: Insights from transgenic mouse models of PyMT-induced breast cancer: recapitulating human breast cancer progression in vivo. Oncogene 2021, 40(3):475–491.

64. Fiaschi-Taesch N, Sicari B, Ubriani K, Cozar-Castellano I, Takane KK, Stewart AF: Mutant parathyroid hormone-related protein, devoid of the nuclear localization signal, markedly inhibits arterial smooth muscle cell cycle and neointima formation by coordinate up-regulation of p15Ink4b and p27kip1. Endocrinology 2009, 150(3):1429–1439.

65. Fiaschi-Taesch N, Takane KK, Masters S, Lopez-Talavera JC, Stewart AF: Parathyroid-hormone-related protein as a regulator of pRb and the cell cycle in arterial smooth muscle. Circulation 2004, 110(2):177–185.

66. Radler PD, Wehde BL, Wagner KU: Crosstalk between STAT5 activation and PI3K/AKT functions in normal and transformed mammary epithelial cells. Mol Cell Endocrinol 2017, 451:31–39.

67. Sotgia F, Schubert W, Pestell RG, Lisanti MP: Genetic ablation of caveolin-1 in mammary epithelial cells increases milk production and hyper-activates STAT5a signaling. Cancer Biol Ther 2006, 5(3):292–297.

68. Dunbar ME, Young P, Zhang JP, McCaughern-Carucci J, Lanske B, Orloff JJ, Karaplis A, Cunha G, Wysolmerski JJ: Stromal cells are critical targets in the regulation of mammary ductal morphogenesis by parathyroid hormone-related protein. Dev Biol 1998, 203(1):75–89.

69. Johnson RW, Sun Y, Ho PWM, Chan ASM, Johnson JA, Pavlos NJ, Sims NA, Martin TJ: Parathyroid Hormone-Related Protein Negatively Regulates Tumor Cell Dormancy Genes in a PTHR1/Cyclic AMP-Independent Manner. Front Endocrinol (Lausanne) 2018, 9:241.

70. Pal B, Chen Y, Vaillant F, Capaldo BD, Joyce R, Song X, Bryant VL, Penington JS, Di Stefano L, Tubau Ribera N, et al. A single-cell RNA expression atlas of normal, preneoplastic and tumorigenic states in the human breast. EMBO J 2021, 40(11):e107333.

71. Liberzon A, Birger C, Thorvaldsdottir H, Ghandi M, Mesirov JP, Tamayo P: The Molecular Signatures Database (MSigDB) hallmark gene set collection. Cell Syst 2015, 1(6):417–425.

72. Fiaschi-Taesch N, Sicari BM, Ubriani K, Bigatel T, Takane KK, Cozar-Castellano I, Bisello A, Law B, Stewart AF: Cellular mechanism through which parathyroid hormone-related protein induces proliferation in arterial smooth muscle cells: definition of an arterial smooth muscle PTHrP/p27kip1 pathway. Circ Res 2006, 99(9):933–942.

73. Gong P, Xia C, Yang Y, Lei W, Yang W, Yu J, Ji Y, Ren L, Ye F: Clinicopathologic profiling and oncologic outcomes of secretory carcinoma of the breast. Sci Rep 2021, 11(1):14738.

74. Hoda RS, Brogi E, Pareja F, Nanjangud G, Murray MP, Weigelt B, Reis-Filho JS, Wen HY: Secretory carcinoma of the breast: clinicopathologic profile of 14 cases emphasising distant metastatic potential. Histopathology 2019, 75(2):213–224.

75. Krings G, Joseph NM, Bean GR, Solomon D, Onodera C, Talevich E, Yeh I, Grenert JP, Hosfield E, Crawford ED, et al. Genomic profiling of breast secretory carcinomas reveals distinct genetics from other breast cancers and similarity to mammary analog secretory carcinomas. Mod Pathol 2017, 30(8):1086–1099.

76. Li Z, Tognon CE, Godinho FJ, Yasaitis L, Hock H, Herschkowitz JI, Lannon CL, Cho E, Kim SJ, Bronson RT, et al. ETV6-NTRK3 fusion oncogene initiates breast cancer from committed mammary progenitors via activation of AP1 complex. Cancer Cell 2007, 12(6):542–558.

77. Strauss BL, Bratthauer GL, Tavassoli FA: STAT 5a expression in the breast is maintained in secretory carcinoma, in contrast to other histologic types. Hum Pathol 2006, 37(5):586–592.

78. Berry JE, Ealba EL, Pettway GJ, Datta NS, Swanson EC, Somerman MJ, McCauley LK: JunB as a downstream mediator of PTHrP actions in cementoblasts. J Bone Miner Res 2006, 21(2):246–257.

79. Ionescu AM, Schwarz EM, Vinson C, Puzas JE, Rosier R, Reynolds PR, O’Keefe RJ: PTHrP modulates chondrocyte differentiation through AP-1 and CREB signaling. J Biol Chem 2001, 276(15):11639–11647.

80. Casimiro S, Guise TA, Chirgwin J: The critical role of the bone microenvironment in cancer metastases. Mol Cell Endocrinol 2009, 310(1-2):71–81.

81. Assaker G, Camirand A, Abdulkarim B, Omeroglu A, Deschenes J, Joseph K, Noman ASM, Ramana Kumar AV, Kremer R, Sabri S: PTHrP, A Biomarker for CNS Metastasis in Triple-Negative Breast Cancer and Selection for Adjuvant Chemotherapy in Node-Negative Disease. JNCI Cancer Spectr 2020, 4(1):pkz063.

82. Skondra M, Gkioka E, Kostakis ID, Pissimissis N, Lembessis P, Pectasides D, Koutsilieris M: Detection of circulating tumor cells in breast cancer patients using multiplex reverse transcription-polymerase chain reaction and specific primers for MGB, PTHRP and KRT19 correlation with clinicopathological features. Anticancer Res 2014, 34(11):6691–6699.

83. Barash I: Stat5 in the mammary gland: controlling normal development and cancer. J Cell Physiol 2006, 209(2):305–313.

84. Ren S, Cai HR, Li M, Furth PA: Loss of Stat5a delays mammary cancer progression in a mouse model. Oncogene 2002, 21(27):4335–4339.

85. Schmidt JW, Wehde BL, Sakamoto K, Triplett AA, Anderson SM, Tsichlis PN, Leone G, Wagner KU: Stat5 regulates the phosphatidylinositol 3-kinase/Akt1 pathway during mammary gland development and tumorigenesis. Mol Cell Biol 2014, 34(7):1363–1377.

86. Nevalainen MT, Xie J, Torhorst J, Bubendorf L, Haas P, Kononen J, Sauter G, Rui H: Signal transducer and activator of transcription-5 activation and breast cancer prognosis. J Clin Oncol 2004, 22(11):2053–2060.

