## Supplementary figures and images for "PTHrP Induces STAT5 Activation, Secretory Differentiation and Mammary Tumor Progression"

### Supplementary Material

#### Additional File 1

A

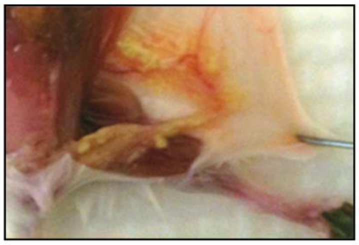

B

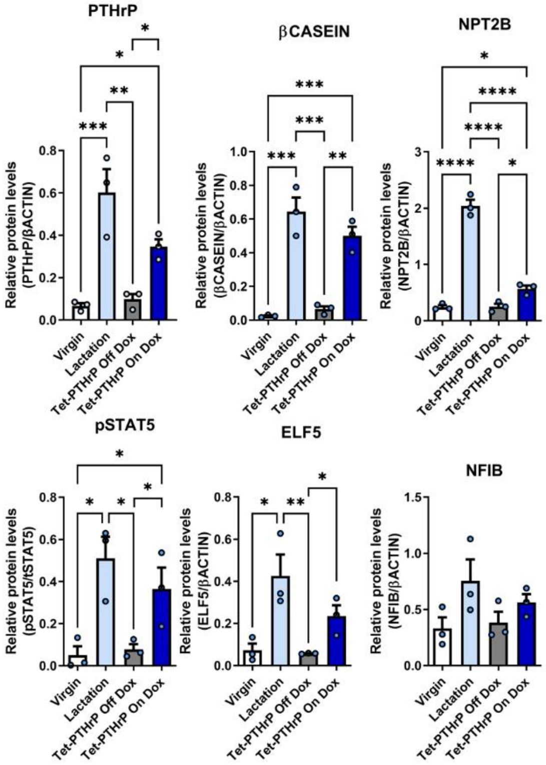

**Additional File 2**

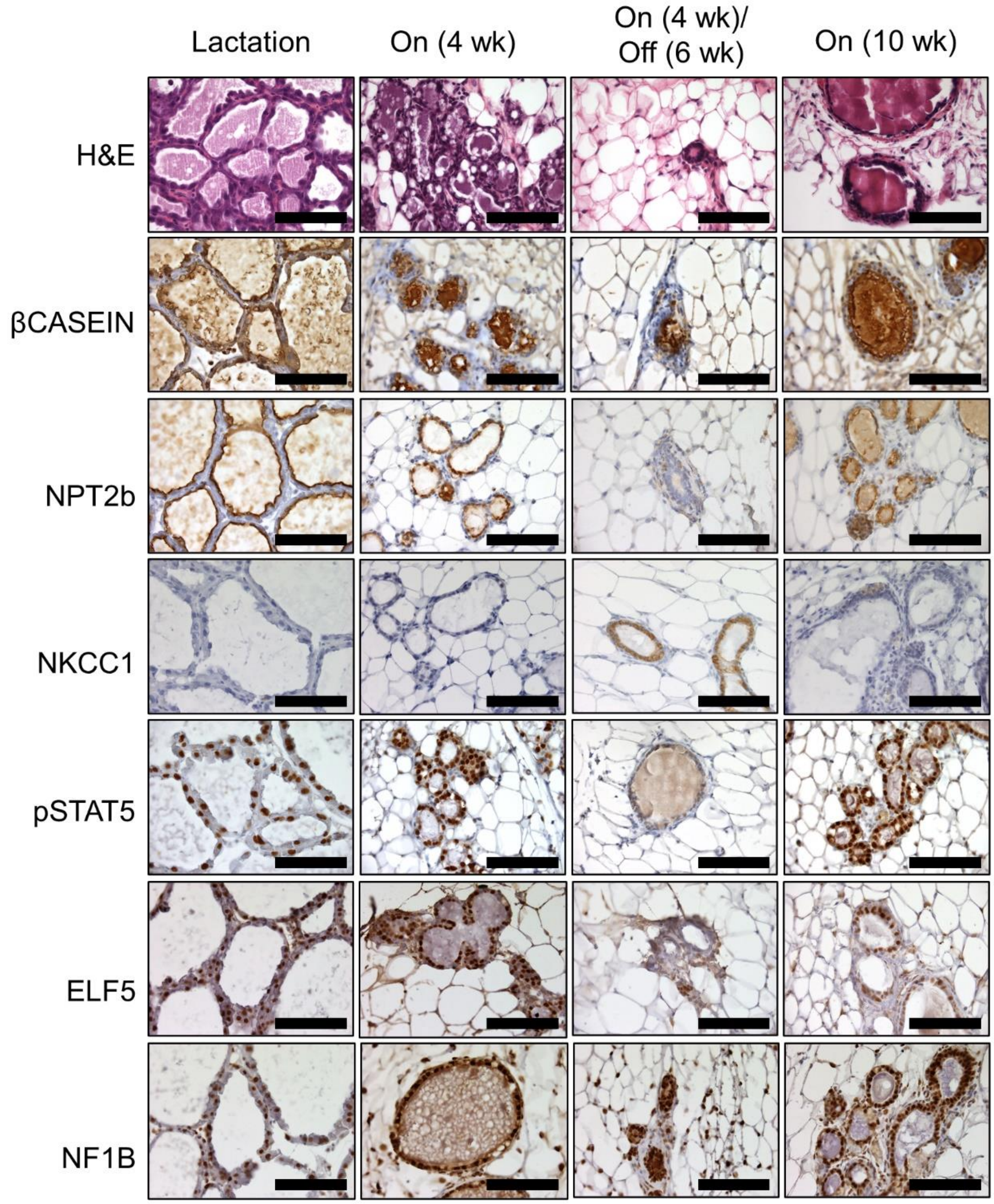

**Additional File 3**

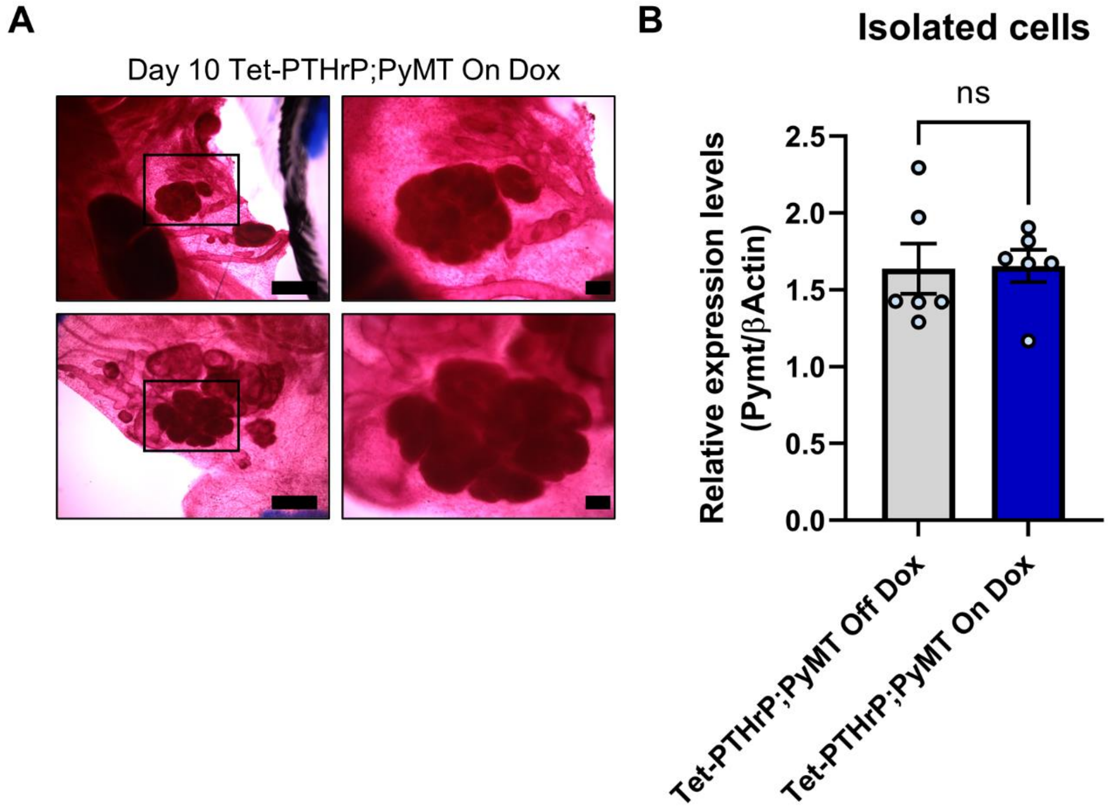

#### Additional File 4

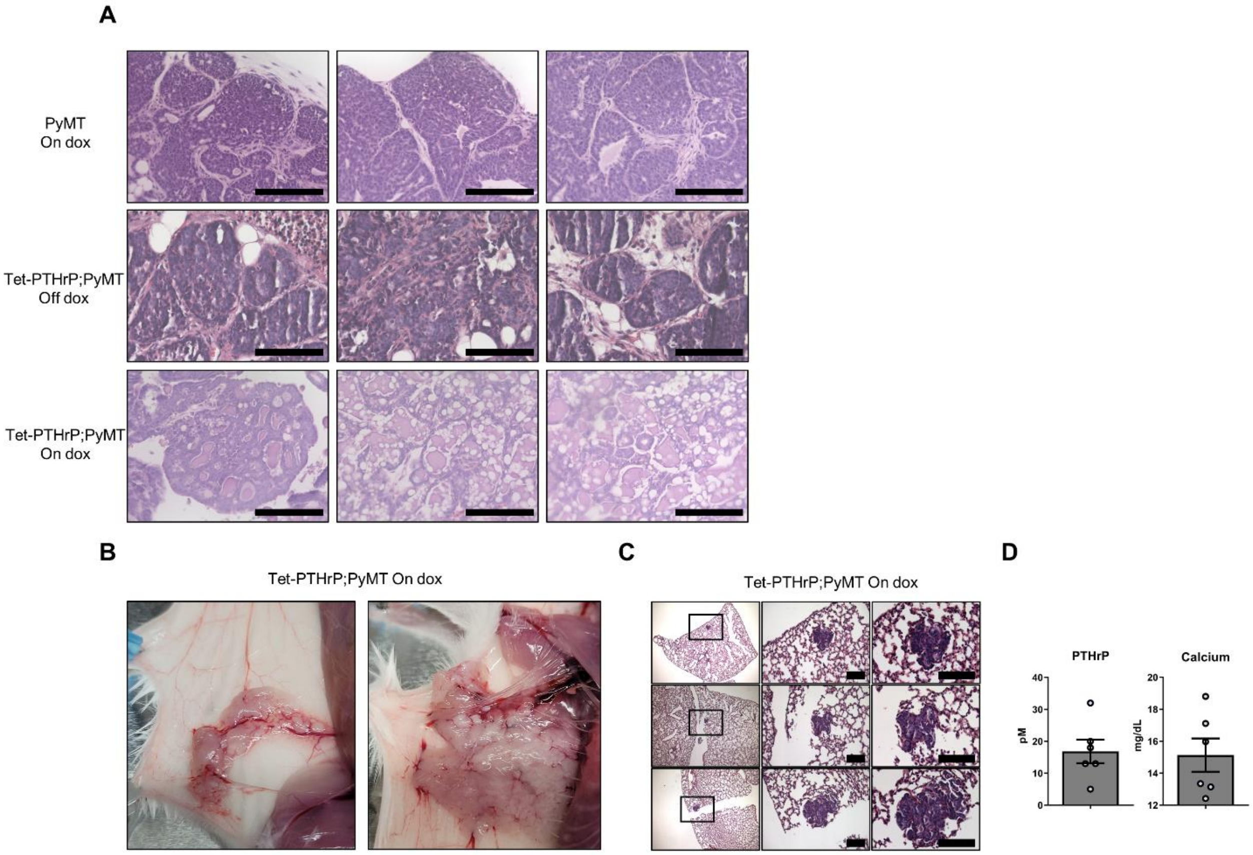

#### Additional File 5

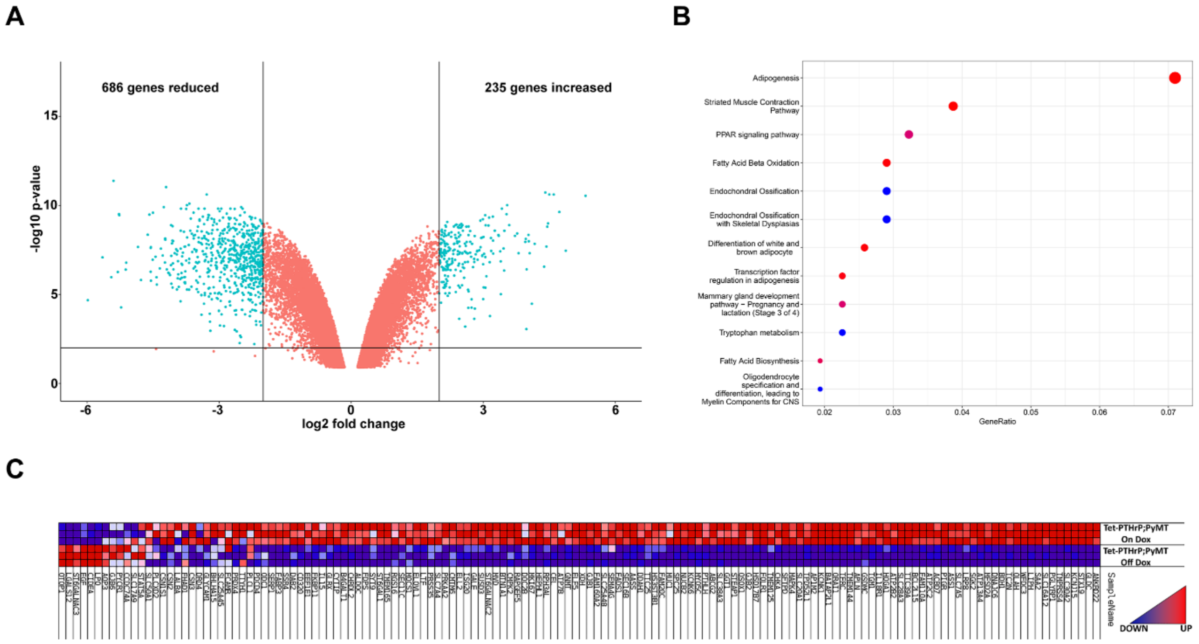
